# Human antiviral protein MxA forms novel metastable membrane-less cytoplasmic condensates exhibiting rapid reversible “crowding”-driven phase transitions

**DOI:** 10.1101/568006

**Authors:** Deodate Davis, Huijuan Yuan, Feng-Xia Liang, Yang-Ming Yang, Jenna Westley, Chris Petzold, Kristen Dancel-Manning, Yan Deng, Joseph Sall, Pravin B. Sehgal

## Abstract

Phase-separated biomolecular condensates of proteins and nucleic acids form functional membrane-less organelles in the mammalian cell cytoplasm and nucleus. We report that the interferon (IFN)-inducible human “myxovirus resistance protein A” (MxA) forms membrane-less metastable condensates in the cytoplasm. Light and electron microscopy studies revealed that transient expression of HA- or GFP-tagged MxA in Huh7, HEK293T or Cos7 cells, or exposure of Huh7 cells to IFN-α2a led to the appearance of MxA in the cytoplasm in membrane-less variably-sized spherical or irregular bodies, in filaments and even a reticulum. 1,6-Hexanediol treatment led to rapid disassembly of these condensates; however, FRAP revealed a relative rigidity with a mobile fraction of only 0.24±0.02 within condensates. In vesicular stomatitis virus (VSV)-infected Huh7 cells, the nucleocapsid (N) protein, which participates in forming phase-separated viral structures, associated with GFP-MxA condensates. Remarkably, the cytoplasmic GFP-MxA condensates disassembled within 1-3 min of exposure of cells to hypotonic medium (40-50 milliosmolar) and reassembled within 0.5-2 min of re-exposure of cells to isotonic medium (310-325 milliosmolar) through multiple cycles. Mechanistically, the extent of cytoplasmic “crowding” regulated this phase-separation process. GFP-MxA condensates also included the DNA sensor protein cyclic GMP-AMP synthase (cGAS), another protein known to be associated with liquid-like condensates. Functionally, GFP-MxA expression inhibited DNA/cGAS-responsive ISG54-luciferase activity but enhanced relative inducibility of ISG54-luc by IFN-α, revealing a physical separation between condensate- and cytosol-based signaling pathways in the cytoplasm. Taken together, the data reveal a new aspect of the cell biology of MxA in the cell cytoplasm.

**Importance:** The human interferon-inducible “myxovirus resistance protein A” (MxA), which displays antiviral activity against several RNA and DNA viruses, exists in the cytoplasm in phase-separated membrane-less metastable condensates of variably-sized spherical or irregular bodies, in filaments and even in a reticulum. MxA condensate formation appeared necessary but not sufficient for antiviral activity. Remarkably, MxA condensates showed the unique property of rapid (within 1-3 min) reversible disassembly and reassembly in intact cells exposed sequentially to hypotonic and isotonic conditions Mechanistically, these phase transitions were regulated by the extent of cytoplasmic “crowding.” Moreover, GFP-MxA condensates included the DNA sensor protein cyclic GMP-AMP synthase (cGAS). Functionally, GFP-MxA expression inhibited DNA/cGAS-responsive ISG54-luciferase activity but enhanced inducibility of ISG54-luc by IFN-α, revealing a biological distinction between condensate- and cytosol-based signaling pathways. Since intracellular edema and ionic changes are hallmarks of cytopathic viral effects, the rapid hypotonicity-driven disassembly of MxA condensates may modulate MxA.function during virus infection.

## Introduction

There is now a growing realization that, in addition to various membrane-bound subcellular compartments, the eukaryotic cell contains organized membrane-less biomolecular condensates (also called supramolecular assemblies) of proteins and nucleic acids which form functional organelles (1–5). Examples include the nucleolus, nuclear speckles, P bodies, stress granules, and several more recent discoveries such as condensates of synapsin or of the DNA sensor protein cyclic GMP-AMP synthase (cGAS) in the cytoplasm or of transcription-associated condensates in the nucleus (1–10). Overall, these condensates have liquid-like properties and are metastable changing to a gel or to filaments commensurate with the cytoplasmic environment (temperature, physical deformation or cytoplasmic “crowding”), and the incorporation of additional proteins, RNA or DNA molecules or posttranslational modifications (1–10). Indeed, DNA and RNA molecules specifically participate in the assembly of such cytoplasmic and nuclear condensates, and in their function (1–10). Recently it has been recognized that replication of negative-strand RNA viruses [e.g. vesicular stomatitis virus (VSV), rabies virus] also occurs in cytoplasmic phase-separated condensates involving viral nucleocapsid (N) with or without additional viral proteins ((11) and citations therein). In the present report we highlight the unexpected discovery that the human interferon (IFN)-inducible “myxovirus resistance protein A” (MxA), which displays antiviral activity against several RNA and DNA viruses (12–14), exists in the cytoplasm in membrane-less metastable condensates.

Human MxA is a cytoplasmic 70-kDa dynamin-family large GTPase which induced in cells exposed to Type I and III IFNs (12–14). There is an extensive literature on the mechanisms of the broad-spectrum antiviral effects of MxA (12–14). These mechanisms include inhibition of viral transcription and translation, and inhibition of the nuclear import of viral nucleic acids (12–14). The antiviral activity of MxA requires the GTPase activity and the ability to form dimers (12–14). MxA has an overall structure consisting of a globular GTPase domain (G domain), a hinge region (BSE region) and a stalk consisting of 4 extended alpha-helices (15–19). Dimerization has been attributed to an interface in the G domain, especially the D250 residue, as well as the stalk region (16, 20) In cell-free studies MxA has been observed to dimerize, tetramerize, as well as self-oligomerize into rods (∼50 nm length) and rings (∼36 nm diameter) based upon temperature and ionic conditions (15–19). Additionally, in cell-free studies, it has been shown that MxA can cause membrane tubulation through a tetra-lysine positively charged loop in the stalk domain (18), but whether this occurs in intact cells is unclear.

Previous investigators reported the occurrence of wild-type MxA in the cytoplasm of mammalian cells in puncta and in various irregularly shaped structures (15–26). GTPase-null mutants which lacked antiviral activity also generated such cytoplasmic structures. Extensive mutational studies revealed that the D250N mutant of MxA, localized to the dimerization interface, remained exclusively dispersed in the cytoplasm and also lacked antiviral activity (20). A prevalent interpretation of studies over the last 15 years has been to suggest that MxA associated with a “subcompartment of the endoplasmic reticulum (ER)” (12-21, 23-26). However, none of the previous studies characterized MxA localization using authentic markers of the ER such as known structural or luminal ER proteins. In our recent studies using reticulon-4 (RTN4), a structural protein of the mammalian ER, as a marker we were unable to localize MxA to the ER (22, 27). We were also puzzled by the diversity of size and shape of cytoplasmic MxA structures in the published MxA literature (15-20, 23-25) which did not appear to match structural features of the ER (22, 27-32).

In this report we summarize an unexpected aspect of the cell biology of MxA structures in the cytoplasm. The present data reveal that cytoplasmic MxA structures were membrane-less condensates exhibiting marked metastability (shape-changing) and rapid and reversible phase transitions in response to cytoplasmic “crowding.” In GFP-MxA expressing VSV-infected cells the viral nucleocapsid (N) protein associated with GFP-MxA condensates concomitant with development of an antiviral phenotype. Moreover, GFP-MxA condensates included the DNA sensor cGAS – a signal-initiating protein also recently shown to form DNA-induced liquid-like condensates in the cell cytoplasm (6). Functionally, GFP-MxA expression inhibited DNA/cGAS-responsive ISG54-luciferase activity but enhanced the relative inducibility of ISG54-luc by IFN-α, revealing a physical separation between condensate- and cytosol-based signaling pathways.

## Results

### Marked variations in shapes and sizes of cytoplasmic MxA structures

A noteworthy feature of the appearance of cytoplasmic MxA structures illustrated in the literature (15-20, 23-25) as well as in our hands (22, 27) was their marked phenotypic variation. Figs. 1A, 1B and 1C provide an overview of the kinds of MxA-positive structures that were observed in HEK293T and Huh7/Huh7.5 cells following transient transfection of the wild-type HA-MxA expression vector with the indicated variations observed even among different cells within the same culture. MxA associated with variably-sized spherical and irregularly shaped structures (Fig. 1A, 1B, 1C; size range: 200-2000 nm), large floret-like structures (arrows in Fig. 1B; size range: 10-20 µm), and a specific crisscross reticular pattern seen in Huh7 or Huh7.5 cells (Fig. 1C, right image, arrow). Endogenous MxA induced in Huh7 cells exposed to IFN-α also formed an irregular cytoplasmic reticulum (Fig. 1D). This MxA reticulum was distinct from the punctate distribution of DLP1/DRP1, another large dynamin-family GTPase which associates with mitochondria (32, 33) (Fig. 1D).

**Fig. 1.**
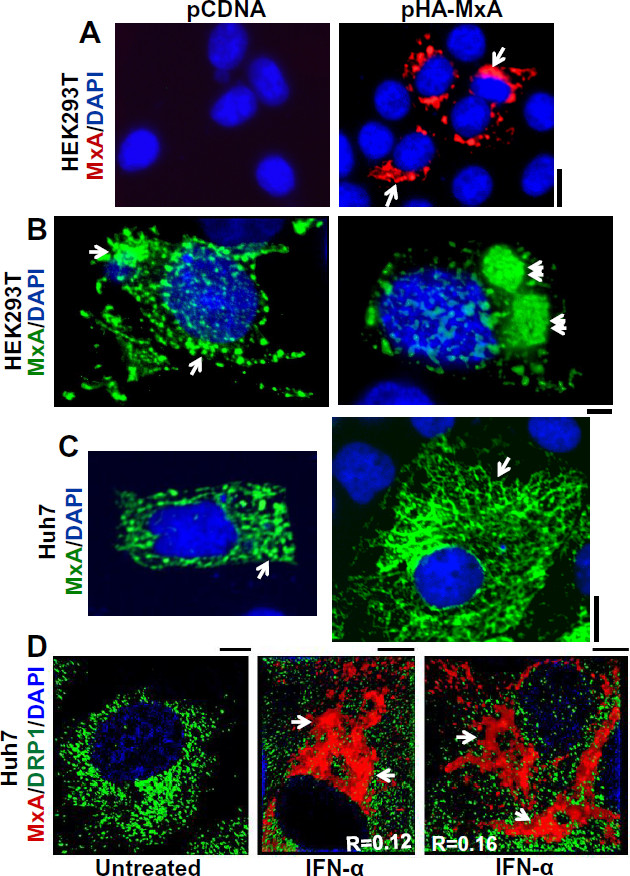
Variation in phenotypes of MxA-positive cytoplasmic structures in different cells. Cultures of respective cells as indicated grown in 35 mm plates or 6-well plates were transiently transfected with pcDNA or the pHA-MxA expression vector, fixed one day later, permeabilized using a digitonin-containing buffer, and the distribution of MxA evaluated using an anti-HA mAb or an anti-MxA rabbit pAb and immunofluorescence methods. Panel A, MxA structures in HEK 293T cells. Imaged using a 40x water immersion objective (scale bar = 20 µm). Panel B, MxA structures in HEK293T cells imaged using an 100x oil immersion objective (scale bar = 10 µm). Arrows point to large compact tubuloreticular MxA-positive structures as observed in 25-35% of transfected cells. Panel C, MxA structures in Huh7 cells imaged using a 40x water immersion objective (scale bar = 10 µm) showing variably-sized bodies (left image) and larger reticular formations in the cytoplasm as observed in 25-35% of transfected cells in the cytoplasm (right image, arrow) (scale bar = 10 µm). Panel D. Huh7 cells were exposed to IFN-α2a (3,000 IU/ml) for 2 days or left untreated, and then fixed and immunostained for endogenous MxA (in red) and endogenous DRP1 (in green) (scale bar = 5 µm).

The phenotypic variation in MxA structures was independent of the protein tag used. Fig. S1 shows that both HA- and GFP-tagged wild-type MxA displayed the same heterogenous structures with complete co-localization of both tagged MxA species in these structures. Critically, the use of GFP-tagged MxA allowed for studies in live cells, including in time-lapse imaging experiments, in correlated fluorescence/light and electron microscopy (CLEM) and analyses of fluorescence recovery after photobleaching (FRAP). In live cells, GFP-MxA formed variably sized spherical structures (Fig. 2A) which showed local movement (see Fig. 5A below), alignment of these along a crisscross pattern (Fig. 2B), as well as formation of an intricate filamentous reticular meshwork (Fig. 2C). As expected from our previous studies (22, 27), this MxA reticulum was distinct from the standard endoplasmic reticulum when tested using an antibody to the ER structural protein RTN4 (28–31) (Fig. 2D).

**Fig. 2.**
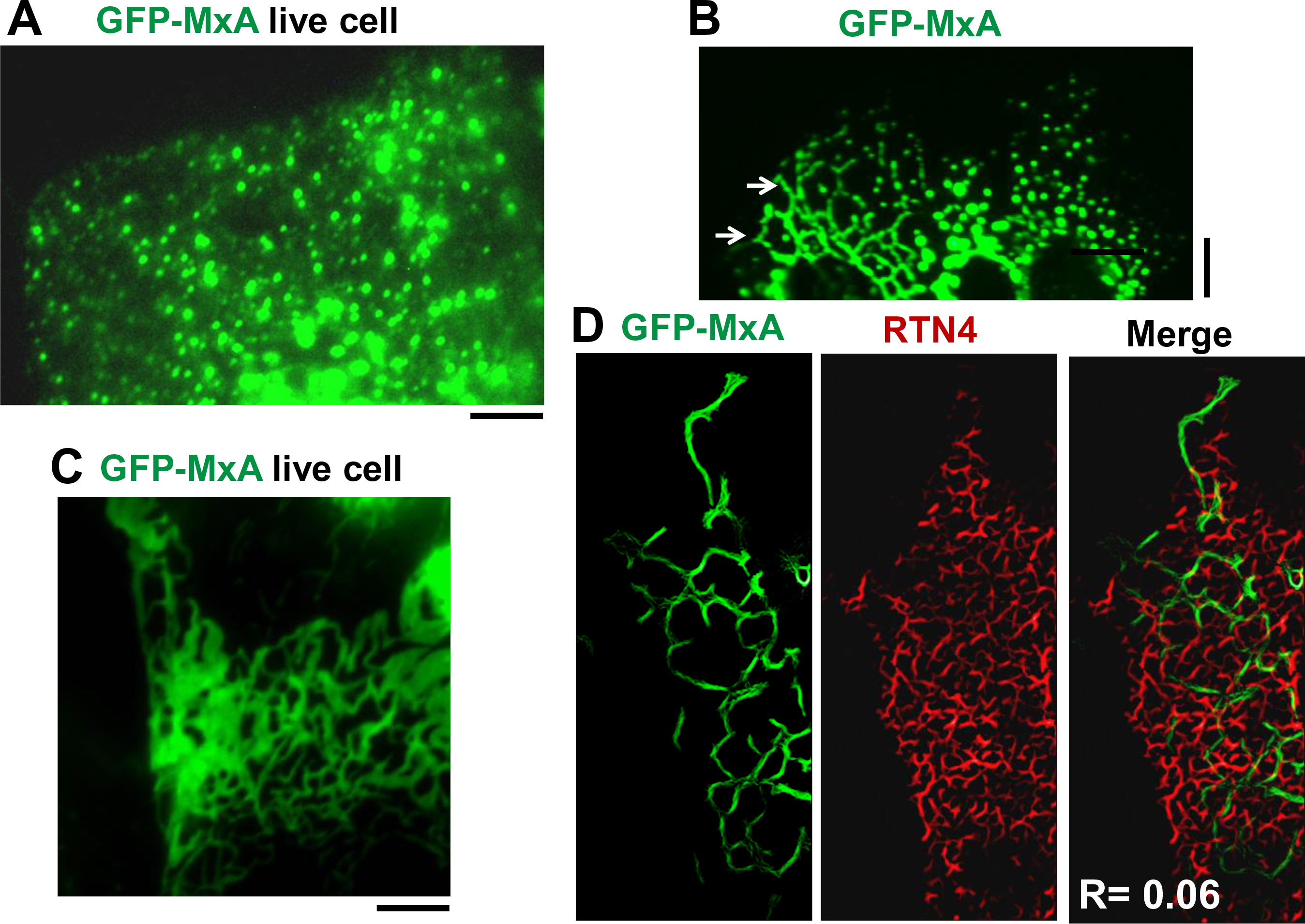
GFP-MxA formed variably-sized cytoplasmic bodies and a reticular meshwork in Huh7 cells distinct from the RTN4-based ER. Huh7 cells in 35 mm plates were transfected with the pGFP-MxA expression vector and the cells imaged before (Panels A and C) or after fixation (Panels B and D). Panel A shows live-cell wide-field 100x oil imaging of the periphery of one cell with GFP-MxA structures of variable size within the same cell one day after transfection; smaller-sized MxA bodies were at the cell periphery with larger structures closer to the cell center. Panel B shows portions of three adjacent cells one day after transfection showing GFP-MxA associated with structures in different configurations, including a reticular pattern of intersecting lines (arrows; left-most cell). Scale bars in Panels A and B = 10 µm. Panel C shows an open meshwork GFP-MxA reticulum in a live Huh7 cells two days after transfection imaged by placing a coverslip on the cells followed by time-lapse imaging using an 100x oil objective. Scale bar = 5 µm. Panel D, Cultures as in Panel C were fixed and then imaged after immunostaining for the ER marker RTN4. The 100x oil images were tubeness filtered using Image J (Fiji). Scale bar = 5 µm. R is the Pearson’s correlation coefficient after automatic Costes’ thresholding.

Figs. S2-S8 summarize a more exhaustive test of the possible localization of MxA to the endoplasmic reticulum using high resolution confocal microscopy and five different markers for the ER (28–31) in both fixed-cell and live-cell labeling studies. These comparisons included HA-MxA and RTN4 (Fig. S2), HA-MxA and HA-atlastin 3 (Figs. S3 and S4), endogenous CLIMP63 as well as mCh-CLIMP63 compared with HA-MxA, GFP-MxA and IFN-α-induced endogenous MxA (Fig. S5), GFP-MxA and RTN4-mCh in live Cos7 and Huh7 cells (Fig. S6), GFP-MxA with mCh-KDEL (Fig. S7), and GFP-MxA with Sec61β-mCh by high resolution confocal microscopy (Fig. S8). As with our previous data (22, 27), the present data did not allow us to localize MxA significantly to any portion of the ER (also see below Fig. S12 for double-label immuno-EM studies). It is noteworthy from the high-resolution confocal data in Fig. S8, that not only are the GFP-MxA structures distinct from Sec61β-mCh but also that the GFP-MxA structures in different cells showed a heterogeneity of shape.

### Cytoplasmic GFP-MxA structures comprise membrane-less condensates

The cytoplasmic GFP-MxA structures were characterized using correlated light and thin-section electron microscopy (CLEM) (34, 35). Fig. S9 illustrates one example of a GFP-MxA expressing cell that was subjected to thin-section EM (TEM). The boxed area in Fig. S9B is shown at higher magnification in Fig. 3A, while boxed area 2 in Fig. S9A is shown at higher magnification in Fig. 3C. Fig. 3A shows that the GFP-MxA structure consisted of electron-dense irregular circular and reticular structures without an external limiting membrane. Similarly, Fig. 3C shows multiple membrane-less condensates of GFP-MxA arranged alongside cytoskeletal elements. As a control, Fig. 3B illustrates clear membrane-containing ER in images derived from cells on the same thin section as Figs. 3A and 3C. Fig. S10 provides a higher magnification image of the central region in Fig. 3C emphasizing the occurrence of GFP-MxA condensates lacking in an external membrane. Single-label immuno-EM studies confirmed the association of MxA with variably-sized small condensates as well as larger heterogenous reticular structures (Fig. S11). Moreover, double-label immuno-EM studies confirmed that the variably-shaped MxA condensates were largely devoid of RTN4 (Fig. S12). Importantly, RTN4 positive structures distinct from MxA were detected in these studies (insets in Figs. S12A and S12B) validating the double-label immuno-EM technique.

**Fig. 3.**
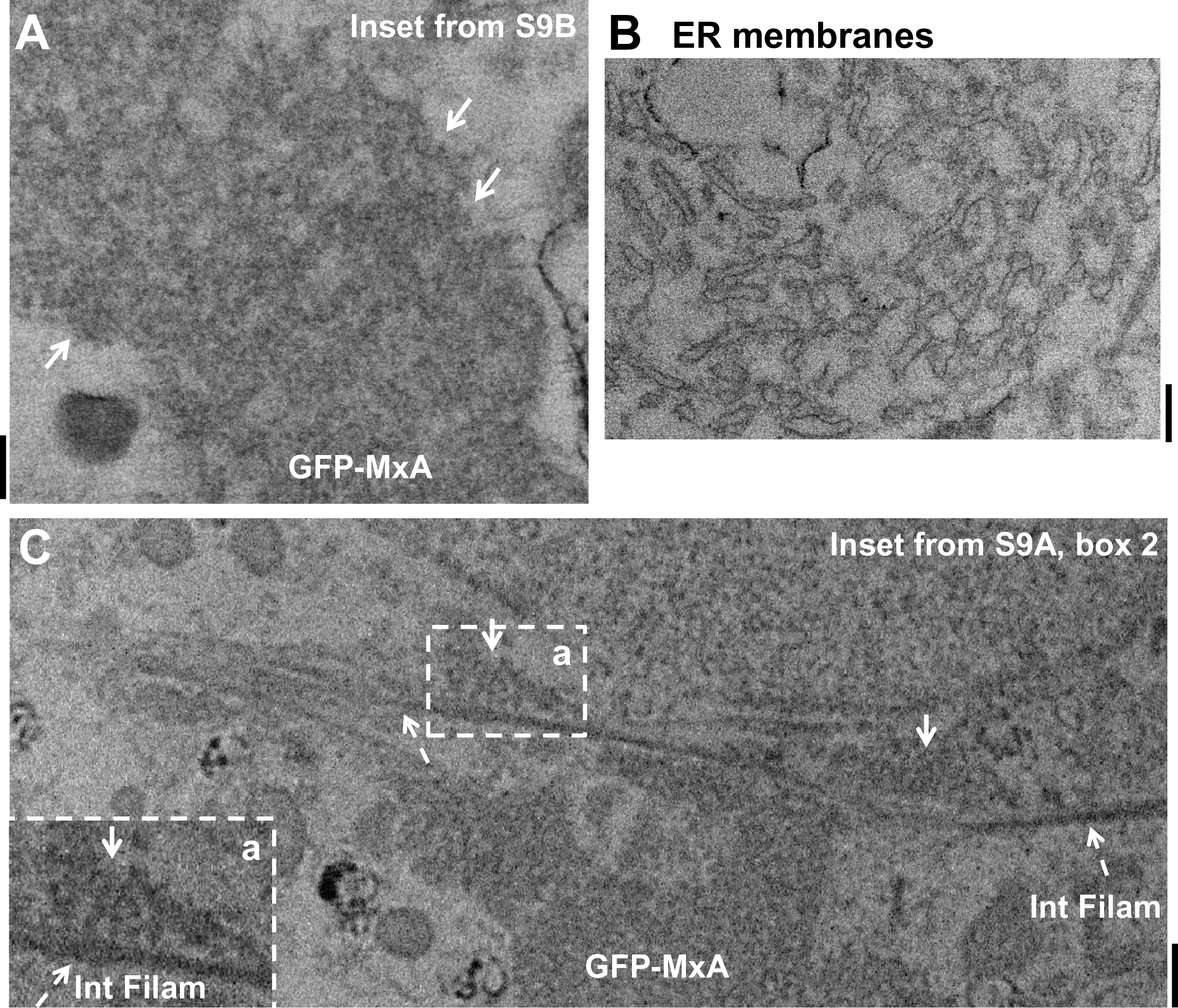
Thin-section EM of GFP-MxA structures in Huh7 cells carried out using a correlated light and electron microscopy (CLEM) approach. Huh7 cells plated sparsely in 35 mm gridded coverslip plates were transiently co-transfected with the pGFP-MxA. Two days later the cultures were fixed with 4% paraformaldehyde for 1 hr at 4°. Confocal imaging was carried out using a tiling protocol to identify the location of specific cells with GFP-MxA structures on the marked grid (Fig. S9). The cultures were then further fixed, embedded and the previously identified grid locations used for serial thin-section EM. The tiled light microscopy data were correlated with the tiled EM data to identify the ultrastructure of the GFP-fluorescent structures (arrows in Fig. S9A) as in the correlated light and electron microscopy (CLEM) procedure. Panel A, higher magnification of thin-section EM image of GFP-MxA structures (arrows) corresponding to inset box in Fig. S9B. Panel B, higher magnification thin-section EM image of a portion of a cell in the same section as in Panel A imaged at the same time, verifying the ability to detect ER membranes in this experiment. Panel C, higher magnification image of thin-section EM of Fig. S9A, inset box 2, showing some of the GFP-MxA structures (arrows) aligned along intermediate filaments (Int Filam; broken arrows). A higher magnification of inset “a” in Panel C is shown in the lower left corner. All scale bars = 200 nm.

### Association of VSV N protein with GFP-MxA condensates

The data in Figs. 4A and 4B confirm the antiviral effect of the expression of GFP-MxA in Huh7 cells by transient transfection. When assayed four hours after VSV infection at a high multiplicity of infection (MOI; >10 pfu/cell) (36), GFP-MxA-positive cells showed a marked reduction in expression of the VSV N protein compared to GFP-MxA-negative cells in the same culture (Figs. 4A and 4B). The higher magnification images in Fig. 4C show a cell from a different region of the same culture as in Figs. 4A and 4B, revealing the co-localization of viral N protein with a subset of GFP-MxA condensates (white single arrows in Fig. 4C). It is noteworthy that there was biochemical heterogeneity in the association between N and GFP-MxA. In Fig. 4C the round dot-like variably-sized condensates showed co-localization (white single arrows) but not the larger irregular GFP-MxA condensates (white double arrows).

**Fig. 4.**
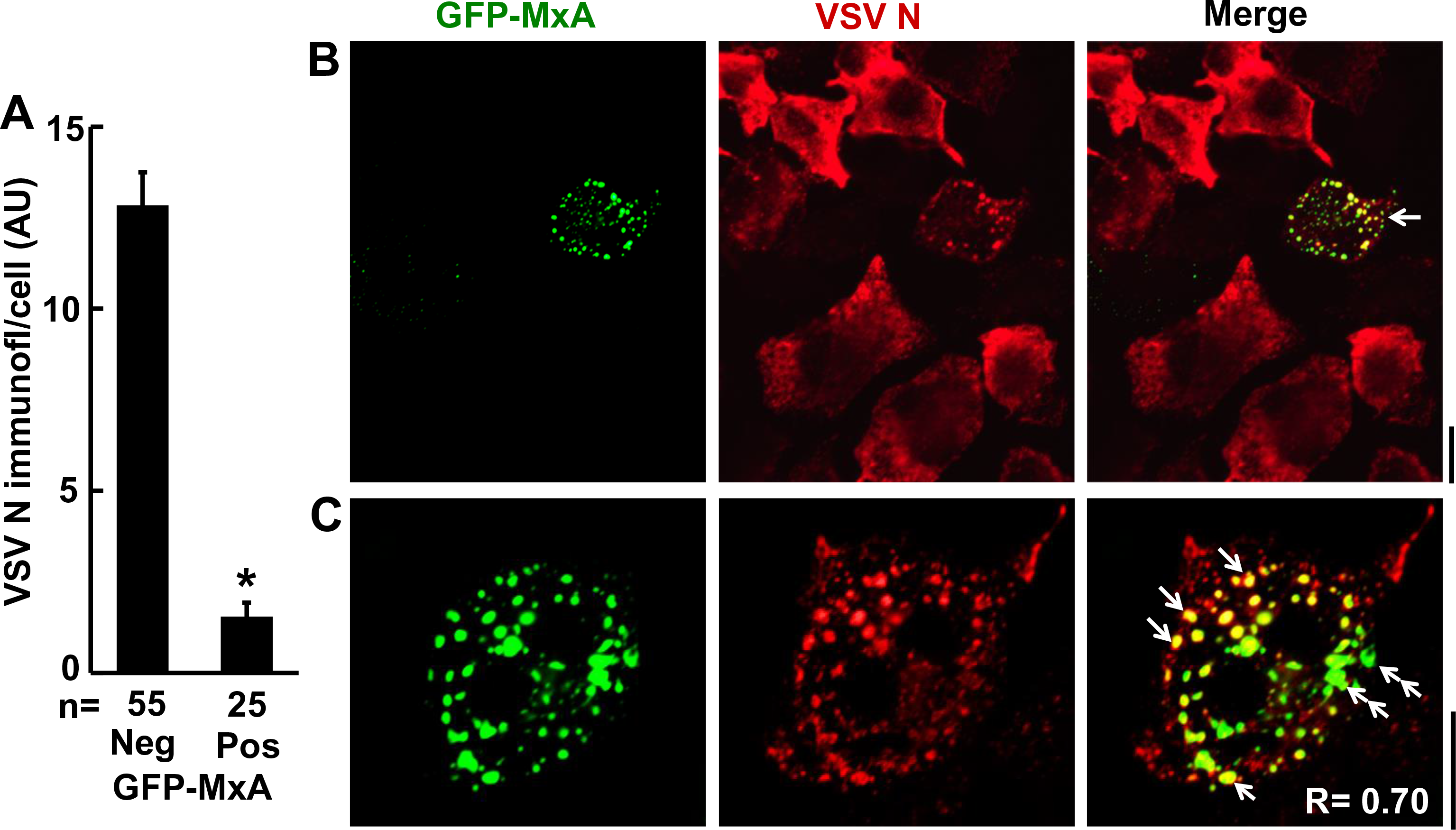
Antiviral effect of MxA: VSV nucleocapsid (N) protein associates with GFP-MxA condensates. Huh7 cells (approx. 2 × 10^5^) per 35 mm plate, transfected with the pGFP-MxA expression vector 1 day earlier, were replenished with 0.25 ml serum-free Eagle’s medium and then 10 µl of a concentrated VSV stock of the wt Orsay strain added (corresponding to MOI >10 pfu/cell). The plates were rocked every 15 min for 1 hr followed by addition of 1 ml of full culture medium. The cultures were fixed at 4 hr after the start of the VSV infection and the extent and localization of N protein expression in individual cells evaluated using immunofluorescence methods (using the mouse anti-N mAb) and Image J for quantitation. Panel A, enumerates N protein expression in GFP-MxA negative and positive cells in the same culture in arbitrary units per cell; * *P* < 0.001. Panel B illustrates a field of GFP-MxA negative cells expressing high levels of N protein surrounding a GFP-MxA-positive cell with limited N protein expression. Panel C shows a higher magnification image of a GFP-MxA positive cells expressing some N protein and the colocalization of the two (white arrows) as well as structures in which the two do not co-localize (white double arrows).

**Fig. 5.**
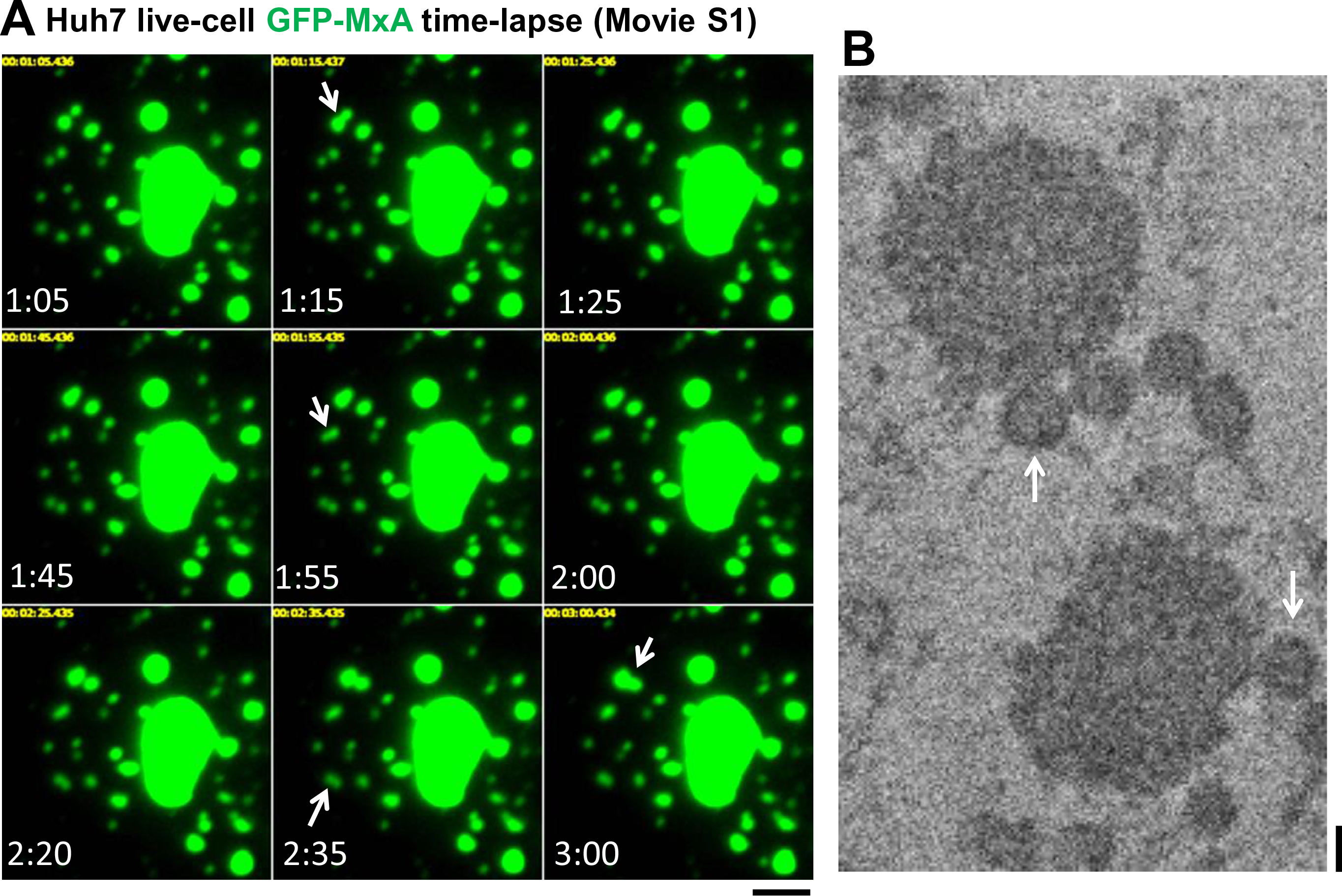
Homotypic association of GFP-MxA bodies to yield larger structures. Panel A. Huh7 cells in 35 mm plates were transfected with the pGFP-MxA expression vector. Two days later the live cells, under a coverslip with a drop of PBS, were imaged using an 100x oil immersion objective and time lapse data collection (images were collected from the periphery of different cells at 5 second intervals for 1-3 min). Figure illustrates images from one time-lapse sequence lasting approximately 3 min (Movie S1) showing four homotypic association events (arrows). The data also show persistence of the combined structure. Scale bar = 5 µm. Panel B. Thin-section EM images from the CLEM data as in Fig. 3 which highlight the homotypic fusion event between small GFP-MxA bodies (arrows) with larger MxA structures all of which lack an external limiting membrane. Scale bar = 200 nm.

### GFP-MxA condensates are metastable and heterogeneous

The GFP-MxA structures were dynamic and changed shape. Live-cell time-lapse imaging revealed that GFP-MxA condensates displayed homotypic fusion. Fig. 5A and Movie S1 show four fusion events within approximately 2 min in the peripheral cytoplasm of a GFP-MxA expressing cell. Fig. 5B shows thin-section EM image of a GFP-MxA expressing cell (from the CLEM experiment in Fig. 3) illustrating small condensates in the vicinity of larger structures, suggestive of an impending fusion event (compare Fig. 5B with Fig. 5A and Movie S1)

The liquid-like nature of the interior of membrane-less condensates is often tested by their rapid disassembly upon exposure of cells to the plasma membrane permeable reagent 1,6-hexanediol and by rapid fluorescence recovery after photo bleaching (FRAP) (3, 37-39). Fig. 6A shows data from an experiment in which hexanediol rapidly disassembled GFP-MxA condensates within 1-2 min. The FRAP analyses in Fig. 6B and C show that the interior of GFP-MxA condensates comprised a mobile fraction of only 0.24 (compared to ∼0.70 for cytoplasmic GFP-STAT3) indicating significant rigidity in MxA condensate structure. This rigidity is consistent with data in Figs. 1, 2 and 3 above (as well as in Fig. 7 below) reporting MxA structures to have multiple odd shapes, be filamentous, comprise reticula both by themselves as well as in association with cellular cytoskeletal elements (Fig. 3B) (22, 27, 40)).

**Fig. 6.**
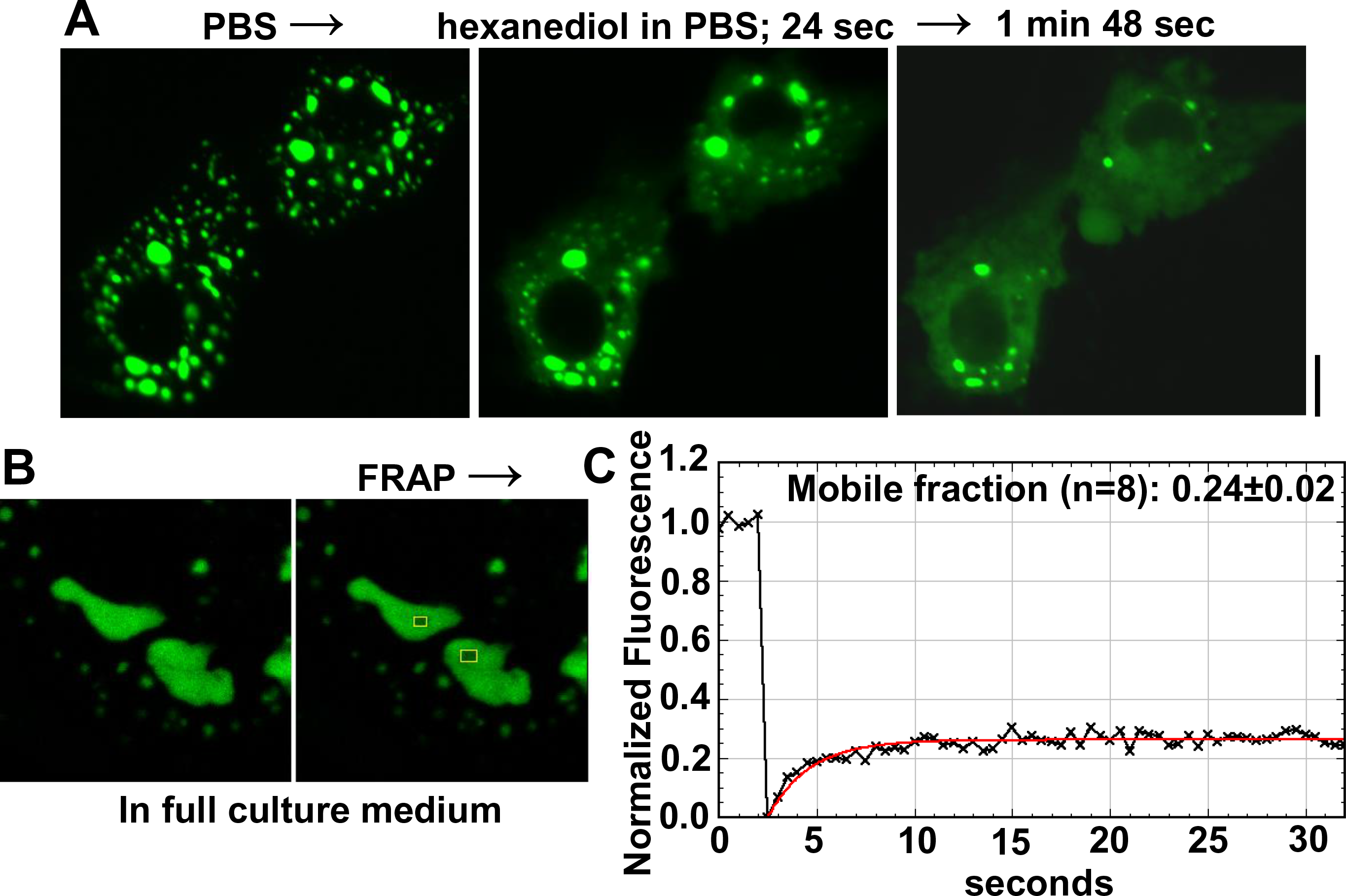
Test of liquid-like properties of GFP-MxA condensates. Panel A, Hexanediol rapidly disassembles GFP-MxA condensates: Huh7 cells expressing GFP-MxA condensates were first imaged in PBS, and then exposed to PBS containing 1,6-hexanediol (5%) and imaged 24 sec and 1 min 48 sec later. Scale bar = 10 µm. Panels B and C. FRAP analyses of replicate (n =8) internal regions of GFP-MxA condensates as summarized in Materials and Methods. Panels B and C: FRAP analyses. Panel B shows the location of two of the bleached spots evaluated. Panel C show the normalized bleaching and recovery plot for one of the spots in Panel B. Overall (n=8), mobile fraction was 0.24 ± 0.02, and t_1/2_ was 2.37 ± 0.35 seconds (mean ± SE). Controls for normalization included unbleached spots in each image, as well as background recording away from the GFP-MxA condensate. As a positive control for mobility, Huh7 cells expressing GFP-STAT3 in the cytoplasm showed a mobile fraction of 0.70 (60, 61).

**Fig. 7.**
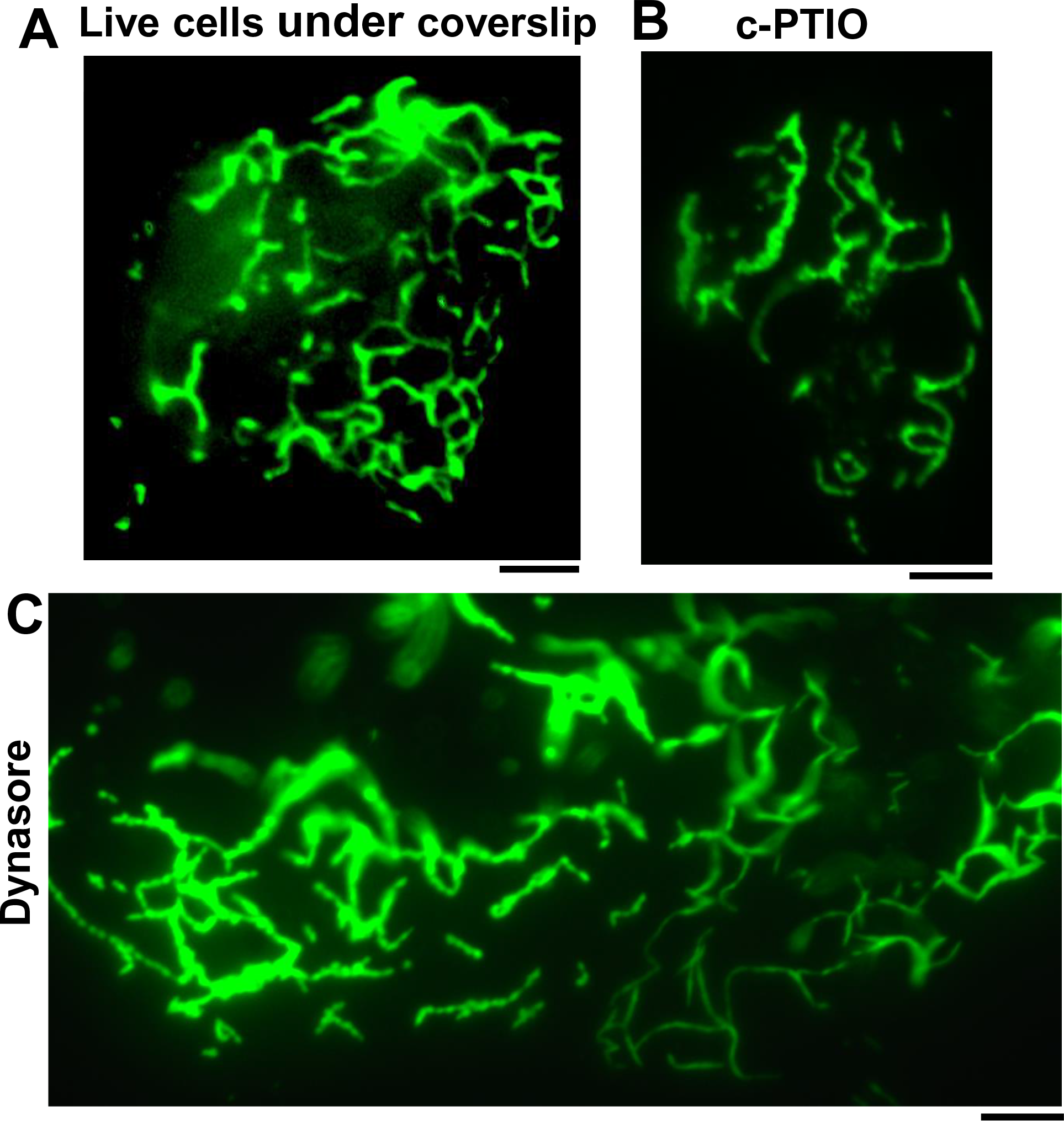
Metastability of GFP-MxA condensates – transition to filamentous reticulum. Panel A, Huh7 cells in 35 mm plates were transfected with the pGFP-MxA expression vector. Two days later the live cells were placed under a coverslip with a drop of PBS, and imaged using an 100x oil immersion objective. By 30-40 min into the imaging session, GFP-MxA was observed to be in a fibrillar meshwork in most cells. Panel B, GFP-MxA expressing cells were treated with the nitric oxide scavenger c-PTIO (300 µM for 7 hrs). Fibrillar conversion of GFP-MxA condensates was first observed using a 40x water immersion objective (not shown) and further confirmed by placing a coverslip and 100x oil imaging with time-lapse data collection. Image shown is one illustrative example of a 100x time-lapse experiment. A time-lapse movie of a different cell is shown in Movie S2. Panel C, GFP-MxA expressing cells were treated with dynasore (20 nM) for 4 days. Fibrillar conversion of GFP-MxA condensates was first observed using a 40x water immersion objective (not shown) and further confirmed by placing a coverslip and 100x oil imaging with time-lapse data collection. Image shown is one illustrative example of a 100x time-lapse experiment. A time-lapse movie of a portion of the same cell Movie S3. All scale bars = 5 µm.

Indeed, shape-changing (“metastability”) is a common feature of membrane-less condensates in the cytoplasm (1–10). When cells expressing the large GFP-MxA condensates were subjected to mechanical deformation for 30-40 min by placing a coverslip on live cells in phosphate-buffered saline (PBS) followed by 100x imaging using an oil-immersion objective, the GFP-MxA condensate often unraveled into an open filamentous meshwork (Fig. 7A). A GFP-MxA filamentous meshwork was also frequently observed in cells in cultures that had been treated with a nitric oxide scavenger 2-4-carboxyphenyl-4,4,5,5-tetramethylimidazoline-1-oxyl-3-oxide (c-PTIO; (41))(Fig. 7B) or with dynasore (42) (Fig. 7C). Movies S2 and S3 illustrate time-lapse imaging of the GFP-MxA reticulum in cells in experiments shown in Figs. 7B and 7C. Filaments in these GFP-MxA reticular meshwork exhibited local oscillatory movements (Movies S2 and S3).

### Rapid and reversible tonicity-driven phase transitions of GFP-MxA condensates

In live-cell studies of GFP-MxA expressing Huh7 cells we made the serendipitous discovery that exposing cells to hypotonic buffer used for swelling cells at the start of many cell fractionation protocols (e.g. erythrocyte lysis buffer, ELB) (40-50 milliosmolar) (43) led to a rapid transition of GFP-MxA from condensates to the cytosol in a matter of 1-3 mins (Fig. 8A and Movie S4). Remarkably, this dispersed cytosolic GFP-MxA could be reassembled back into condensates (but different from the original ones in the same cell) by exposing cells to buffer made isotonic using sucrose (addition of 0.275 M sucrose to ELB) or regular RPMI 1640 medium (Fig. 8A and Movie S5; in particular Movie S4 shows the same cell through disassembly and then reassembly of GFP-MxA condensates). GFP-MxA condensates could be repeatedly cycled back and forth through disassembly and reassembly multiple times (Fig. 8A shows 2 cycles; we have carried out up to 3 cycles). Sequential exposure of GFP-MxA expressing cells to ELB supplemented with different concentrations of sucrose identified the hypotonic range between 80-120 milliosmolar for when disassembly became apparent (data not shown). The majority (80-90%) of GFP-MxA expressing cells in a culture showed the phase transitions of disassembly and reassembly irrespective of the level of expression of MxA in individual cells (low, medium or high levels in small puncta or large condensates). Exposing GFP-MxA-expressing cells to hypertonic buffers (approximately 500 millimolar) also resulted in a change of shape from spherical MxA bodies to elongated cigar-shaped structures; this shape change was reversible upon returning the cells to isotonic medium and even disassembly upon shift to hypotonic buffer (ELB).

**Fig. 8.**
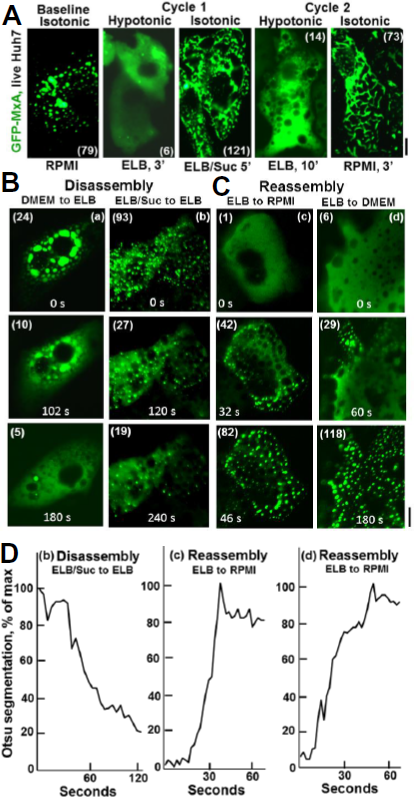
Rapid and reversible tonicity-driven disassembly and reassembly of GFP-MxA condensates in the cytoplasm of Huh7 cells. Panel A, GFP-MxA expressing cultures 2 days after transfection were sequentially imaged on a warm stage (37°C) using a 40x water immersion objective under isotonic conditions (RPMI 1640 medium; “RPMI”), switched to hypotonic medium (erythrocyte lysis buffer, “ELB”, approximately 40 milliosmolar) and imaged 3 min later, then switched to isotonic buffer (ELB supplemented with 0.275 sucrose, net approximately 325 milliosmolar) and imaged 5 min later, followed by a further cycle of hypotonic and isotonic media as indicated. Scale bar = 10 µm. Numbers in parenthesis are the separate “objects” counted in each using Otsu segmentation (Image J) and automatic investigator-independent thresholding. Scale bar = 10 µm. At the end of this experiment the cultures were fixed and immunostained for the ER marker RTN4 to confirm that the reassembled GFP-MxA condensates were independent of the ER (data shown in Fig. S13B). Movie S4 shows time-lapse images (5 sec apart) of the first cycle of disassembly and then reassembly of GFP-MxA condensates in the same cell different experiment. Panels B and C, show illustrative examples (labelled a, b, c, and d) of disassembly and reassembly of GFP-MxA condensates taken from time-lapse imaging experiments as generated by the medium/buffer changes indicated. Numbers in parenthesis are the number of objects counted in each image using Otsu segmentation protocol. Scale bar = 10 µm. Panel D, Quantitation of the speed of disassembly and reassembly using Otsu segmentation metrics in time-lapse movies corresponding to examples b, c and d in Panel B. Movie S5 shows the reassembly cell (c) in Fig. 8C and 8D.

Time-lapse imaging of cells subjected to disassembly and reassembly showed these phase transitions to be rapid processes initiated within 30 seconds and completed by 1-3 mins (Fig. 8B and 8C; Movies S4 and S5). Disassembly was typically completed by 2-3 minutes and reassembly a bit faster by 1-2 min. As a control, cells expressing the N1-GFP tag alone did not show any condensates, nor any assembly and disassembly phase transitions (Fig. S13A). Additionally, GFP-MxA condensates taken through two cycles of disassembly and reassembly (cells at the conclusion of the experiment shown in Fig. 8A), were again confirmed to be distinct from the RTN4-positive ER (Fig. S13B).

### Mechanism of GFP-MxA condensate formation and disassembly: regulation by cytoplasmic crowding

The observation that GFP-MxA condensates rapidly disassembled upon hypotonic cell swelling and rapidly reassembled upon isotonic cell recovery (albeit into different structures in the same cell) (Fig. 8) suggested that the mechanism of cytoplasmic crowding (4, 8) might be involved in the regulation of GFP-MxA phase separation. The observation that disassembly and reassembly took place in similar manner in cells expressing high, modest or trace levels of GFP-MxA (experiments such as in Fig. 8) suggested that the critical issue was not the concentration of GFP-MxA in the cytoplasm but the state of dilution of other cytoplasmic contents. The data in Fig. 9A provide important insight into the occurrence of this mechanism during condensate reassembly. When swollen cells with hypotonically disassembled GFP-MxA condensates were switched to isotonic PBS, there was a microscopically visible phase transition in the cytoplasm in ∼15-30 seconds evidenced by accumulation of dark GFP-free droplets (white solid arrows in 33 and 37 sec panels in Fig. 9A) which crowded GFP-MxA into the inter-droplet regions (white dotted arrows in 33 and 37 sec panels in Fig. 9A). Over the next several seconds, the GFP-MxA crowded into the inter-droplet regions formed condensations with the persistence and enlargement of the dark GFP-free droplets (white dotted and solid arrows respectively in the 52 and 72 sec panels in Fig. 9A). The data in Fig. 9A, top row, provide direct imaging evidence of a phase transition in the cytoplasm that occurred upon switching to isotonic medium and subsequent GFP-MxA formation in the inter-droplet regions.

**Fig. 9.**
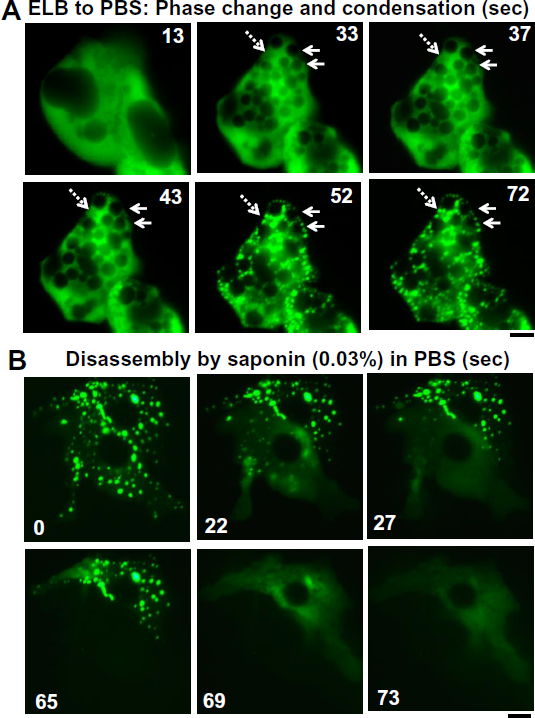
Cytoplasmic crowding regulates assembly and disassembly of GFP-MxA condensates. Panel A, Huh7 cells expressing GFP-MxA condensates were switched to hypotonic ELB as in Fig. 8A, left side, and GFP-MxA disassembly imaged. The culture was then switched to isotonic PBS during continuous repeated imaging over the next 2 min. Panel A shows a montage of 6 images of the same cell from just prior to and for approximately 1 min after switching to isotonic PBS. White arrows show a rapid phase transition characterized by dark droplets crowding the GFP-MxA into the inter-droplet region. White double arrows point to GFP-MxA condensation in the inter-droplet regions. Panel B, Disassembly of GFP-MxA condensates in the presence of saponin in isotonic buffer. The same cells containing GFP-MxA condensates were imaged repeatedly from just prior to and for approximately 1 min after switching to saponin (0.03%)-containing PBS. Scale bars = 10 µm.

The time-lapse Movie S5 provides a clear example of this phase transition in the cytoplasm upon switching to isotonic buffer – first the appearance of dark droplets, and then condensation of GFP-MxA in the inter-droplet regions. These data are consistent with the regulation of GFP-MxA condensation by a cytoplasmic crowding mechanism.

Fig. 9B provides further evidence from the disassembly point of view in support of the cytoplasmic crowding mechanism. In the montage in Fig. 9B, cells containing GFP-MxA condensates were exposed to isotonic PBS containing saponin (0.03%) which is known to permeabilize the plasma membrane ((43) and citations therein). The data in Fig. 9B show that within seconds there was marked disassembly of GFP-MxA condensates with loss of GFP-MxA into the culture medium. That permeabilizing the plasma membrane led to rapid disassembly of GFP-MxA condensates is consistent with the regulation of GFP-MxA condensate structures by a cytoplasmic crowding mechanism. The observation that hypotonic cell swelling rapidly disassembled GFP-MxA condensates (Fig. 8) is also consistent with the regulation of GFP-MxA condensate structure by a cytoplasmic crowding mechanism. Taken together, the data in Figs. 8 and 9, and Movies S4 and S5, suggest that cytoplasmic crowding (Alberti, 2017) regulated the balance between the pool of GFP-MxA in condensates and that in the cytosol.

### GFP-MxA condensates physically segregate cytoplasmic signaling pathways

Recently, because (a) cGAS, a DNA sensor protein which enhanced a pathway stimulating IFN-α and –β production, was observed to be in a liquid-like cytoplasmic condensate (6), and (b) MxA has been shown to enhance IFN-α and –β production in Huh7.5 cells (44) we investigated whether cGAS might co-localize with GFP-MxA condensates. Fig. 10A, left column, and Fig. S14 show that GFP-MxA condensates did indeed include cGAS. Moreover, Fig. 10A, right column, shows that GFP-MxA and cGAS significantly reassembled together after one cycle of hypotonic disassembly and isotonic reassembly suggesting a level of specificity in this association.

**Fig. 10.**
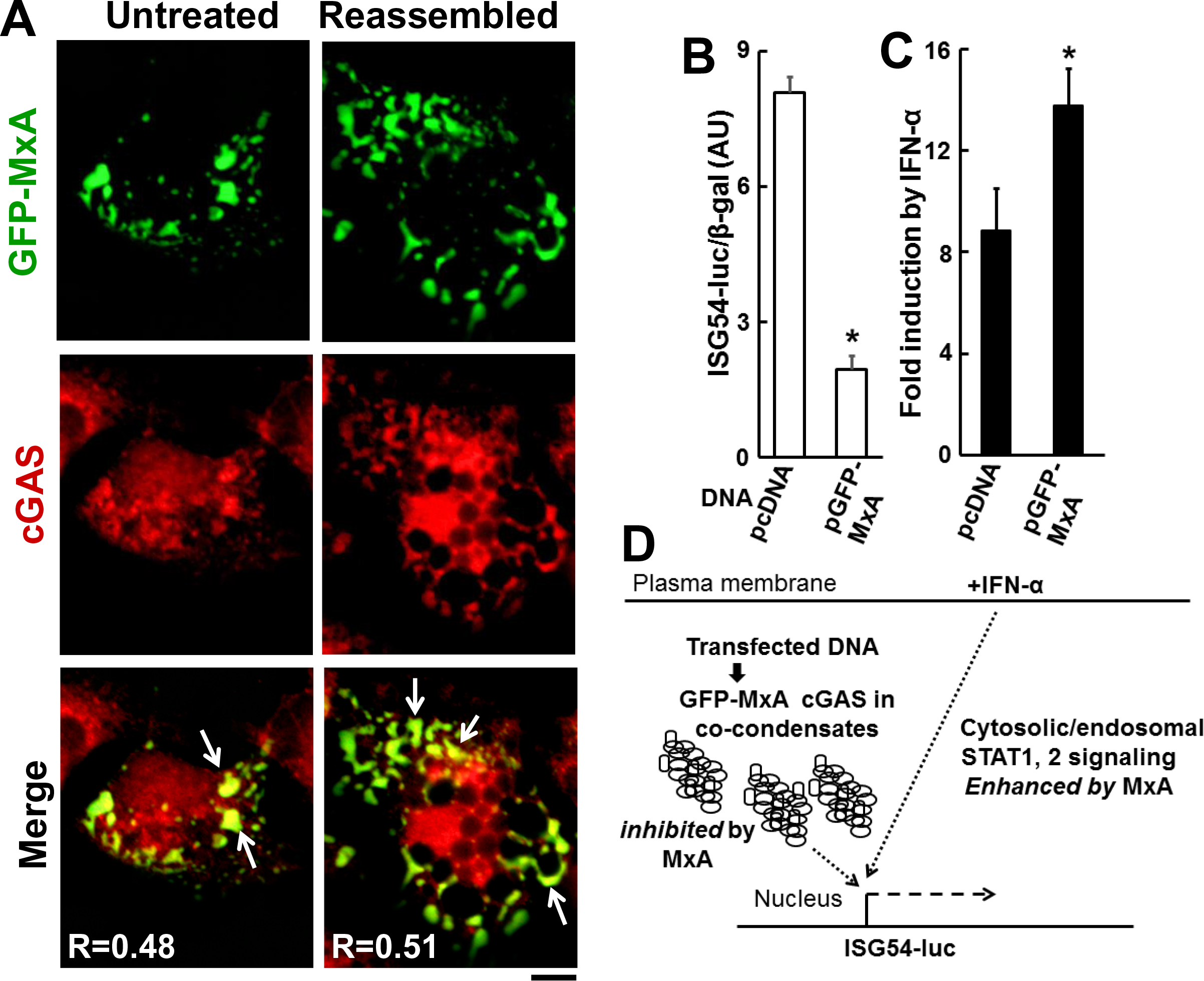
Association of cGAS with GFP-MxA condensates and physical segregation of signaling pathways through the cytoplasm. Panel A, GFP-MxA expressing cultures 2 days after transfection were either left untreated in PBS or cycled through hypotonic disassembly and isotonic reassembly (same cultures shown, in part, in Fig. 9A) followed by fixation and evaluation of cGAS colocalization with GFP-MxA by immunofluorescence imaging. R is the Pearson’s correlation coefficient after automatic Costes’ thresholding. Scale bar = 10 µm. Panel B, Huh7 cells in 6-well plates were transfected with DNA mixtures containing ISG54-luc construct (1 µg/well), pCH110 (constitutive β-galactosidase reporter; 1 µg/well) together with either pcDNA (4 µg/well) or pGFP-MxA DNA (4 µg/well) with each variable evaluated in triplicate. Seventy-two hrs after transfection, GFP-MxA expressing cells were confirmed to show condensate formation by fluorescence microscopy, and then all cultures harvested for respective reporter assays. ISG54 luciferase activity data were normalized to β-galactosidase activity in each sample and are expressed in arbitrary units (AU) (mean ±SE). * *P* <0.05. Panel C, in an additional experiment similar to that outlined in panel B, respective cultures (in triplicate per variable) were exposed to IFN-α (2,5000 IU/ml) for the last 18 hr of the 3-day post-transfection period. Normalized ISG54-luc activity was first derived as in Panel B, and then the fold-induction by IFN-α derived by expressing data from the IFN-α-treated cultures relative to those from respective IFN-free cultures (mean ± SE); * *P* <0.05. Panel D, Physical segregation of cytoplasmic MxA/ DNA/cGAS condensate-associated and IFN-α cytosol/endosomal-associated signaling pathways.

The data showing the inclusion of cGAS in GFP-MxA condensates (Figs. 10A and S14) led us to investigate whether GFP-MxA might functionally affect the cGAS signaling pathway. The reporter construct ISG54-luc corresponding to the promoter regulatory elements of the interferon-induced gene 54 (ISG54) responds to activation of the DNA/cGAS/STING pathway in the cytoplasm (6). Additionally, the ISG54-luc reporter also responds to stimulation of cells by IFN-α at the plasma membrane through activation of a cytosol/endosome-associated STAT1/STAT2 pathway (45, 46). Parenthetically, MxA has been previously shown to stimulate endosome-associated IL-6, BMP4 and BMP9 transcriptional signaling (22, 43). Thus, we first used the ISG54-luc reporter to investigate the effect of GFP-MxA expression in cells into which pcDNA or pGFP-MxA DNA had been introduced by transient co-transfection followed 3 days later by verification of condensate formation in GFP-MxA expressing cells by fluorescence microscopy, and then harvesting all cultures for respective reporter assays (inducible luciferase and the constitutive control β-galactosidase activities). The data in Fig. 10B show a marked (>5-fold) inhibition of the ISG54-luciferase activity in cells expressing GFP-MxA. We then evaluated the ability of IFN-α to stimulate ISG54-luciferase activity in cells transfected with pcDNA or pGFP-MxA DNA three days earlier by exposing these to IFN-α for the last 18 hrs of the experiment. The data in Fig. 10C show that, as expected, IFN-α stimulated the ISG54-luc reporter in terms of fold-inducibility (∼8-fold). Remarkably, this IFN-α-stimulated induction was enhanced further in the presence of GFP-MxA (Fig. 10C). Taken together, the data in Figs. 10B and 10C suggest that GFP-MxA/cGAS co-condensates may comprise a locale for functional inhibition of the DNA/cGAS/ISG54-luc pathway by MxA physically segregated from cytosolic/endosomal IFN-α/STAT1,2/ISG54-luc pathway which is stimulated by cytosolic MxA (Fig. 10D).

## Discussion

Regulated phase transitions of biomolecular condensates in the cytoplasm and nucleus are increasingly viewed as critical in diverse cellular functions (1–10). However, the rules governing these phase transitions remain largely undiscovered. We report the discovery that cytoplasmic MxA condensates underwent rapid and reversible bulk disassembly and then reassembly, each largely completed in a matter of less than 1-3 minutes, in cells cycled through exposure to hypotonic and then isotonic buffers. The rapidity and reversibility of these transitions through multiple cycles were exceptionally striking. This discovery appears similar to “cytoplasmic crowding-based” phase transitions observed by Delarue et al (8) using synthetic multimeric nanoparticle reporters in yeast cells grown overnight in hypertonic medium or in the cytoplasm of HEK293T cells treated overnight with rapamycin to enhance ribosome crowding. The distinctive feature of the MxA phase transitions observed in the present study is the rapidity of the observed changes: bulk disassembly within 1-3 min of the start of hypotonic swelling, and bulk reassembly within 0.5-2 min of return to isotonic medium. Mechanistically, the live-cell imaging evidence showed that the process of GFP-MxA recondensation in isotonic buffer (after hypotonic disassembly) was preceded by a phase separation in the cytoplasm within 15-30 seconds comprising the accumulation of dark GFP-free (water) droplets which crowded GFP-MxA into inter-droplet regions where subsequent condensation was seeded. Conversely, permeabilization of the plasma membrane using saponin led to rapid disassembly of the GFP-MxA condensates. The data support a model in which cytoplasmic crowding regulated the formation and maintenance of the structure of GFP-MxA condensates.

Previous cell-free studies had shown that recombinant wild-type MxA at high concentrations readily assembled into dimers, tetramers, and higher-order oligomers forming rods (∼50 nm length) and rings (∼36 nm diameter) based upon temperature and ionic conditions and associated with and caused membrane tubulation (15-19, 21, 22). This led to the interpretation that the variably sized and shaped MxA structures represented some “subcompartment of the endoplasmic reticulum” but without any ER-specific data provided in the literature (15–19). Our previous studies (22, 27), and the data in Figs. S2-S8 provide evidence for the lack of significant association of MxA with the ER. In contrast, we provide evidence in the present study for the occurrence of MxA in membrane-less variably-sized and variably-shaped metastable condensates in the cytoplasm of living cells. We reinterpret MxA mutational studies reporting the phenotype of MxA structures in the cytoplasm of human HEK293T, Huh7, Vero and Cos7 cells to show that the GTPase activity was not required for the formation of cytoplasmic condensates (20). Mutants which lacked GTPase activity and lacked antiviral activity formed large cytoplasmic condensates (20). However, the D250N mutant, which renders the mutant GFP-MxA protein cytosolic, also lacked antiviral activity (20). Taken together, a reinterpretation of the previously published data suggests that the ability of MxA to form condensates may be required for but may not be sufficient for the manifestation of antiviral activity.

As for how MxA condensates might contribute to an antiviral mechanism, data from several previous studies have emphasized the sequestration of viral nucleocapsid proteins critical to viral replication in cytoplasmic MxA structures (12-14, 17, 20, 23-25). We note specifically that Fig. 2 in Kochs et al (2002) (23) showed the co-association of MxA and N protein of La Crosse virus by thin-section EM and by immuno-EM in membrane-less perinuclear structures that did not associate with any cytoplasmic membrane compartment. These authors, already in 2002, went on to show that MxA cross-associated with the N proteins of La Crosse, Rift Valley fever and Bunyamvera viruses in cross-immunoprecipitation assays. It is only now that we realize that the N proteins of such viruses is included in a liquid-like phase-separated compartment (see Heinrich et al 2018 and citations therein). The present data in Fig. 4 showing the association of The VSV N protein with GFP-MxA condensates, shown to be membrane-less in Fig. 3, confirm the observations of Kochs et al (2002) (23).

In as much as intracellular edema and ionic changes are hallmarks of cytopathic viral infections (47–51), the rapid hypotonicity-driven disassembly of MxA condensates may function to disable the antiviral activity of MxA. Indeed, in a manner anticipating the recent report of altered phase transitions of synthetic reporter protein oligomers in cells with cytoplasmic “crowding” by ribosomes (8), Montasir et al in their studies of vaccinia pneumonia already in 1966 commented that “the most prominent early cytopathic change was intracellular edema evidenced by low electron density to the background cytoplasmic material and dilatation of endoplasmic reticulum” (51). Reduced cytoplasmic crowding in swollen cells exposed to hypotonic buffers would mimic the intracellular edema in virus-infected cells reported by Montasir et al (51), triggering a phase transition (disassembly) of MxA condensates resulting in a reduction in MxA antiviral activity. Experimentally, whether edema-like hypotonic disassembly of MxA condensates affects its antiviral activity remains unclear.

The GFP-MxA condensates in live cells were metastable and changed shape. One the one hand, GFP-MxA condensates were rapidly disassembled by hexanediol suggesting liquid-like internal properties. On the other hand, FRAP analyses showed only a 24% mobile fraction in the interior of the GFP-MxA condensates. This relative rigidity of GFP-MxA structures was keeping with the ability of MxA structures to form odd-shaped structures, filaments, and even extensive reticular networks. We observed homotypic fusion events, as well as a change in shape to filamentous meshworks upon mechanical deformation, exposure to nitric oxide scavenging for several hours and even treatment of cells with the dynamin-family inhibitor dynasore (for 4 days). Parenthetically, although dynasore was reported not to inhibit MxA GTPase activity “significantly” in cell-free assays lasting 20-30 min (42), the published data refer only to a short-term cell-free assay, and even these data do show a small decrease in MxA GTPase activity in the presence of dynasore (42). Whether cellular proteins in addition to MxA contribute to GFP-MxA condensate structure remains unclear.

Du and Chen have previously reported that the DNA sensor cGAS formed liquid-like condensates (6). We observed that GFP-MxA condensates included included cGAS. These two proteins re-associated following one cycle of disassembly and reassembly suggesting specificity in this interaction (Fig. 10A). Because (a) DNA-activated cGAS condensates stimulate the cGAS/STING pathway which culminates in activation of the ISG54-luc reporter and IFN-α and –β production (6 and citations therein), and (b) Shi et al reported recently that transient expression of MxA in Huh7.4 cells led to expression of IFN-α and –β (44), we investigated the effect of MxA on ISG54-luc activity in cells transfected with DNA. In comparison with pcDNA-transfected cultures, pGFP-MxA DNA transfection (and, thus, GFP-MxA expression and co-condensate formation with cGAS) markedly inhibited ISG54-luc activity as assayed 3 days after DNA introduction into cells (Fig. 10B). However, exposure to IFN-α elicited a strong stimulation of the ISG54-luc reporter in both pcDNA or pGFP-MxA DNA transfected cultures (Fig. 10C); the stimulation was higher in the presence of GFP-MxA. The enhancement by MxA of IFN-α signaling transiting through cytosolic/endosomal STAT1, 2 pathways (45, 46) recapitulates the stimulation of by MxA of endosome-associated transcriptional by the cytokines IL-6, BMP4 and BMP9 (22, 43). An interpretation of the contrasting data in Fig. 10B and 10C on the effects of GFP-MxA would be that MxA in co-condensates with cGAS inhibited a cGAS-condensate-based pathway but enhanced the cytosol/endosome-based IFN-α pathway to the same transcriptional reporter (Fig. 10D). Thus, more generally, the formation of condensates by signal regulatory proteins may serve to physically segregate signaling pathways within the cytoplasm.

To summarize, the present studies shed light on a new aspect of the cell biology of MxA in the cytoplasm – the formation of metastable membrane-less condensates which can undergo rapid and reversible phase transitions. MxA condensates include cGAS and appear to functionally affect the DNA/cGAS signaling pathway. The juxtaposition of the previous MxA mutational literature with the present observations suggests that the ability to form cytoplasmic condensates is necessary for but not sufficient for antiviral activity. The present data identify human MxA as a protein which generates cytoplasmic condensates subject to cytoplasmic crowding-driven disassembly and reassembly in Huh7 cells on the time scale of 1-3 minutes. Functionally, the present studies highlight a novel example of the physical segregation of condensate-based and cytosol-based signaling pathways in the cytoplasm.

## Materials and Methods

### Cells and cell culture

Human kidney cancer cell line HEK293T was obtained from the American Type Culture Collection (ATCC). Human hepatoma cell lines Huh7 and its derivative Huh7.5 (52) were gifts from Dr. Charles M. Rice, The Rockefeller University. Cos7 cells were a gift from Dr. Koko Murakami, New York Medical College. The respective cell lines were grown in DMEM supplemented with 10% v/v fetal bovine serum (FBS) in 90 mm plates. For experiments Huh7/7.5 cells were grown in regular 6-well plates or 35 mm dishes, while HEK293T cells were grown in similar plates coated with fibronectin, collagen and bovine serum albumin (respectively 1 µg/ml, 30 µg/ml and 10 µg/ml in coating medium) (53). For CLEM, Huh7 cells were grown sparsely in 35 mm gridded 1.5 mm coverslip plates (Cat. No. P35G-1.5-14-C-GRID; MatTek Corporation, Ashland, MA). Recombinant human IFN-α2a was purchased from BioVision (Milpitas, CA). In the present experiments Huh7 cultures were exposed for two days to IFN-α at a concentration of 3,000 IU/ml prior in DMEM supplemented with 2% FBS prior to prior to fixation and immunofluorescence analyses (22).

### Plasmids and transient transfection

The HA-tagged human MxA expression vector (cloned into a pcDNA3 vector) was a gift from Dr. Otto Haller (University of Freiburg, Germany) (12, 43) and was the exact same MxA expression construct used by Stertz et al (25) with the HA tag located on the N-terminal side of the MxA coding sequence. The GFP (1-248)-tagged human MxA expression vector was a gift of Dr. Jovan Pavlovic (University of Zurich, Switzerland) (54, 55); the GFP tag was located on the N-terminal side of the MxA coding sequence. HA-tagged ATL3 vector was a gift from Dr. Craig Blackstone (National Institutes of Health) (56, 57) and the mCherry-tagged CLIMP63, RTN4, Sec61β and KDEL expression vectors were gifts from Dr. Jason E. Lee and Gia Voeltz (University of Colorado at Boulder) (29). The interferon responsive luciferase reporter constructs ISRE-luc (Stratagene) and ISG54-luc were gifts of Dr. David E. Levy (Dept. of Pathology, New York University School of Medicine). Transient transfections were carried out using just subconfluent cultures in 35 mm plates or in wells of a 6-well plate using DNA in the range of 0.3-2 µg/culture and the Polyfect reagent (Qiagen, Germantown, MD) and the manufacturer’s protocol. Transient transfections for luciferase reporter activity were carried out in 6-well plates, included the β-galactosidase reporter pCH110, and luciferase activity in the cell extracts normalized for β-galactosidase expression were quantitated as summarized earlier (22).

### VSV stock and virus infection

A stock of the wild-type Orsay strain of VSV (titer: 9 × 10^8^ pfu/ml) was a gift of Dr. Douglas S. Lyles (Dept. of Biochemistry, Wake Forest School of Medicine, Winston-Salem, NC). Virus infection was carried out essentially as described by Carey et al (2008). Briefly, Huh7 cells (approx. 2 × 10^5^) per 35 mm plate, previously transfected with the pGFP-MxA expression vector (1-2 days earlier), were replenished with 0.25 ml serum-free Eagle’s medium and 10 µl of the concentrated VSV stock added (corresponding to MOI >10 pfu/cell). The plates were rocked every 15 min for 1 hr followed by addition of 1 ml of full culture medium. For the experiment shown in Fig. 4, the cultures were fixed at 4 hr after the start of the VSV infection.

### Immunofluorescence imaging

Typically, 1-2 days after transient transfection of respective vectors, the cultures were fixed using cold paraformaldehyde (4%) for 1 hr and then permeabilized using a buffer containing <u>digitonin</u> (50 µg/ml)/sucrose (0.3M) (22, 41, 53, 57, 58). Single-label and double-label immunofluorescence assays were carried out using antibodies as indicated in SI, with the double-label assays performed sequentially (22, 41, 53, 57, 58). Fluorescence was imaged as previously reported (22, 41, 53, 57, 58) using an erect Zeiss AxioImager M2 motorized microscopy system with Zeiss W N-Achroplan 40X/NA0.75 water immersion or Zeiss EC Plan-Neofluor 100X/NA1.3 oil objectives equipped with an high-resolution RGB HRc AxioCam camera and AxioVision 4.8.1 software in a 1388 × 1040 pixel high speed color capture mode. Images in z-stacks were captured using Zeiss AxioImager software; these stacks were then deconvolved and rendered in 3-D using the 64-bit version of the Zeiss AxioVision software. Deconvolution and tubeness filtering of 2-D images was carried out using Image J (Fiji) software. Colocalization analyses were carried out using Image J software (Fiji) and deriving the Pearson’s colocalization coefficient R with Costes’ automatic thresholding (59). Line scans were carried out using AxioVision 4.8.1 software.

High-resolution immunofluorescence/fluorescence imaging of selected cultures was carried out using a Leica SP5 II confocal microscopy system using a Leica HCX APO L 40x/0.8 NA water immersion objective or a Zeiss Confocal 880 Airyscan system (100x oil/xx NA objective).

Live-cell imaging of GFP-MxA and mCherry-tagged ER markers was carried out in cells grown in 35 mm plates using the upright the Zeiss AxioImager 2 equipped with a warm (37°) stage and a 40x water immersion objective, and also by placing a coverslip on the sheet of live cells and imaging using the 100x oil objective (as above) with data capture in a time-lapse or z-stack mode (using Axiovision 4.8.1 software).

FRAP experiments were performed on Zeiss LSM880 confocal microscope with Zeiss Plan-Apochromat 63x/1.4 Oil objective on randomly picked 9 cells. Five pre-bleach images were acquired before 25 cycles of bleaching with full power of Argon 488nm laser on small area (∼3 μm^2)^ which reduces intensity to ∼50%. Fluorescence recovery was monitored continuously for 30 seconds at 2 frames/sec. All image series were aligned and registered with Fiji ImageJ’s SIFT plugin and FRAP analyses of MF and t1/2 were performed afterwards with a Jython script (https://imagej.net/Analyze_FRAP_movies_with_a_Jython_script) incorporated into Image J (Fiji).

### Phase transition experiments

Live GFP-MxA expressing cells in 35 mm plates were imaged using a 40x water-immersion objective 2-5 days after transient transfection in growth medium or serum-free RPMI 1640 medium or in phosphate-buffered saline (PBS). After collecting baseline images of MxA condensates (including time-lapse sequences), the isotonic buffers were removed using a Pasteur pipette and replaced with hypotonic ELB (10 mM NaCl, 10 mM Tris, pH 7.4, 3 mM MgCl_2_) followed by time-lapse imaging at 2-5 second intervals. After 5-10 min (by which time >85% of cells showed complete disassembly of the GFP-MxA condensates), the cultures were subsequently sequentially exposed to additional various isotonic and hypotonic buffers as enumerated in the respective figure legends.

### Electron microscopy

For immuno-EM, the cultures were fixed with paraformaldehyde (4%) for 20 min at 37°C, and then another 40 min at room temperature (53). After fixation, the cells were washed with ice-cold PBS, scraped into an Eppendorf tube and pelleted by centrifugation for 15 sec. The cell pellets were continued to be fixed in freshly made 2% paraformaldehyde in PBS containing 0.2% glutaraldehyde, pH 7.2-7.4 for 4 hours at 4°C. After washing with PBS, the cells were embedded with 10% gelatin, infused with sucrose, and cryo-sectioned at 80 nm thickness onto 200 mesh carbon-formvar coated copper grids. For single MxA labeling, the grids were blocked with 1% coldwater fish skin gelatin (Sigma) for 5 min, incubated with MxA antibody (rabbit pAb) in blocking solution for 2 hours at room temperature. Following washing with PBS, gold-conjugated secondary antibodies (15 nm Protein A-gold, Cell Microscopy Center, University Medical Center Utrecht, 35584 CX Utrecht, The Netherlands, or 18 nm Colloidal Gold-AffiniPure Goat Anti-Rabbit IgG, Jackson ImmunoResearch Laboratories, Inc., West Grove, PA) were applied for 1 hr. The grids were fixed in 1% glutaraldehyde for 5 min, washed with distilled water, contrasted and embedded in a mixture of 3% uranyl acetate and 2% methylcellulose in a ratio of 1 to 9. For double immunolabeling, goat anti-RTN4 antibody was applied first, and the 6 nm colloidal gold-conjugated donkey anti-goat IgG (Jackson ImmunoResearch Laboratories, Inc., West Grove, PA) as secondary antibody. After fixing with 1% glutaraldehyde for 10 min, rabbit anti-MxA antibody were applied, and 15 nm protein A gold was used as the secondary. Electron microscopy was carried out using a Philips CM-12 electron microscope (FEI; Eindhoven, The Netherlands) and images photographed with using a Gatan (4K × 2.7K) digital camera (Gatan, Inc., Pleasanton, CA).

For CLEM (34, 35), Huh7 cells plated sparsely in 35 mm gridded coverslip plates were transiently transfected with the pGFP-MxA vector. Two days later the cultures were fixed with 4% paraformaldehyde for 1 hr at 4°. Confocal imaging was carried out using a tiling protocol to identify the location of specific cells with GFP-MxA structures on the marked grid. The cultures were then further fixed (2.5% glutaraldehyde for 2 hours at 4°C, post-fixed with 1% osmium tetroxide for 1.5 hours at room temperature), and embedded (in EMbed 812; Electron Microscopy Sciences, Hatfield, PA). The previously identified grid locations were used for serial thin-sectioning (60 nm), mounted on 200 mesh thin bar copper grids and stained with uranyl acetate and lead citrate using standard methods. The tiled light microscopy data were correlated with the tiled EM thin-section data to identify the ultrastructure of the GFP-fluorescent structures.

### Antibody reagents

Rabbit pAb to human MxA (also referred to as human Mx1) (H-285) (sc-50509), goat pAb to RTN4/NogoB (N18) (sc-11027) and mouse mAbs to cGAS (D-9)(sc-515777), β-tubulin (2-28-33) (sc-23949) and vimentin (V9) (sc-6260) were purchased from Santa Cruz Biotechnology Inc. (Santa Cruz, CA). Rabbit pAbs to atlastin 3 (ATL3) (ab104262) and to giantin (1-469 fragment) (ab24586) were purchased from Abcam Inc. (Cambridge, MA). Mouse mAb to the HA tag (262K, #2362) was purchased from Cell Signaling (Dancers, MA), while that to the VSV nucleocapsid (N) designated 10G4 was a gift from Dr. Douglas S. Lyles (Wake Forest School of Medicine, NC). Respective AlexaFluor 488- and AlexaFluor 594-tagged secondary donkey antibodies to rabbit (A-11008 and A-11012), mouse (A-21202 and A-21203) or goat (A-11055 and A-11058) IgG were from Invitrogen Molecular Probes (Eugene, OR).

## ACKNOWLEDGMENTS

Supported in part by funding from the New York Medical College, and personal funding from PBS. We thank Drs. Joseph D. Etlinger and Kenneth Lerea for insightful discussions.

## Author contributions

>DD, PBS, HY and FL designed the studies. DD, HY, JW, YMY and PBS carried out the cell culture experiments, collected all the wide-field microscopy data including live-cell imaging, and also analyzed all the data (wide-field, confocal and EM). YD carried out all the high-resolution confocal imaging and FRAP. FL, CP, KD-M and JS carried out all of the electron microscopy, including CLEM, and collected the EM data. PBS and DD compiled the data figures and PBS wrote the manuscript. All authors read and approved the final manuscript.

## Legends to Supplementary Figures

**Fig. S1.**
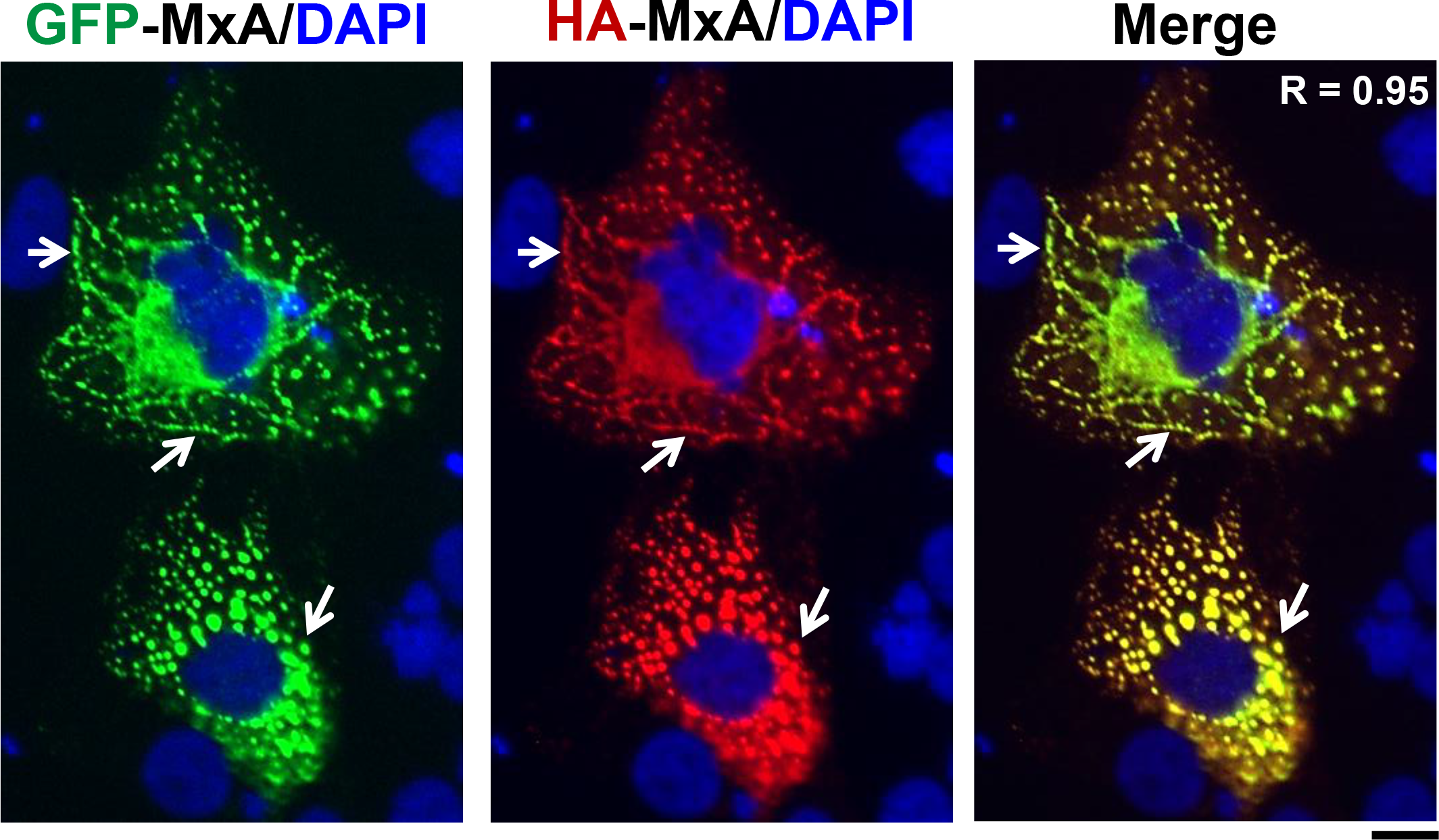
Coincidence of subcellular localization of GFP-MxA with HA-MxA in variably-sized and shaped cytoplasmic structures in Huh7 cells. Cells were transiently transfected with a 1:1 mixture of expression constructs for GFP-MxA and HA-MxA. Two days later the cells were fixed, immunostained using the anti-HA mAb in red to display HA-MxA. Data show coincidence of HA-MxA (in red) with GFP-MxA (in green) structures irrespective of their variation in size and shape (arrows). Scale bar = 10 µm. R = 0.95 indicated in the merged image corresponds to the Pearson’s R coefficient after automatic Costes’ thresholding.

**Fig. S2.**
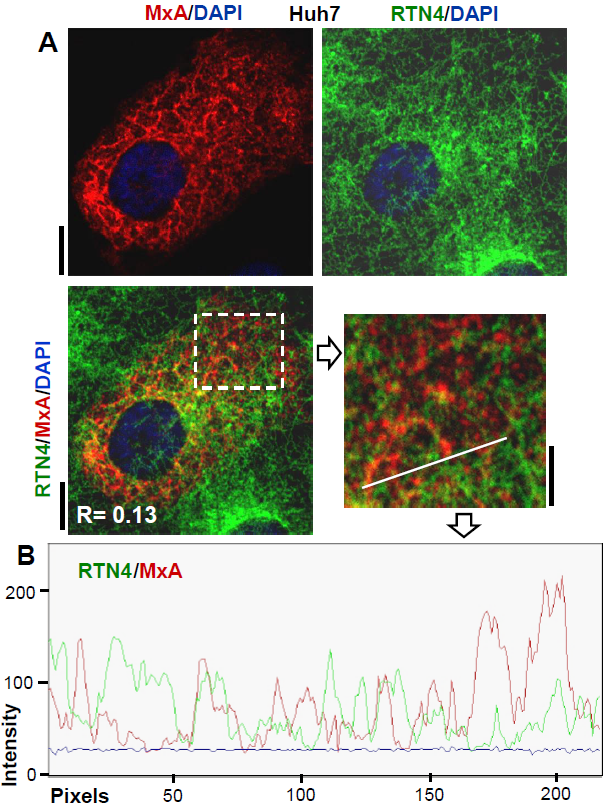
Confocal imaging of the MxA-reticulum in Huh7 cells interspersed alongside but distinct from the standard RTN4-based ER. Double-label immunofluorescence analyses were carried as indicated in the figure after transient transfection of Huh7 cells in 35 mm plates with the HA-MxA expression vector. An anti-RTN4 goat pAb was used to display RTN4 (in green), and an anti-HA mAb was used to display HA-MxA (in red) together with DAPI in sequential staining. Immunofluorescence imaging was carried out using confocal microscopy with a 40x water immersion objective. Panel A illustrates imaging data from a representative cell corresponding to 25-35% of cells in the culture (scale bars on left side = 10 µm). The indicated inset is shown at higher magnification (scale bars on right side = 5 µm). White line in the inset indicate the line scan depicted in Panel B. R value indicated in the merged image in Panel A corresponds to the respective Pearson’s R coefficients (after automatic Costes’ thresholding).

**Fig. S3.**
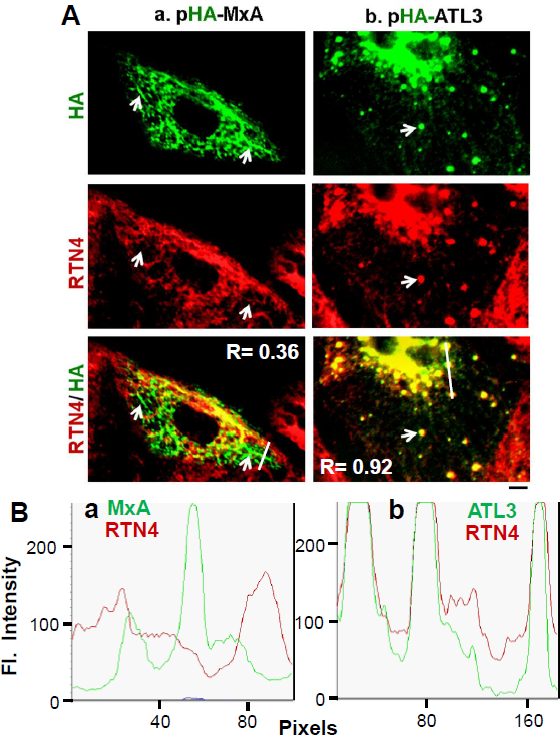
MxA-positive cytoplasmic structures in Huh7.5 cells were distinct from the standard RTN4- and ATL3-based ER (40x imaging). Huh7.5 cells were transfected with vectors for HA-tagged MxA (Panel Aa) or HA-tagged ATL3 (Panel Ab) as indicated, the cultures (in 35 mm plates) were fixed one day later and the distribution of the HA-tag and RTN4 sequentially evaluated using the mouse anti-HA mAb and then a goat anti-RTN4 pAb. Images were collected using a 40x water immersion objective. In Panel Aa arrows in the HA-MxA panels point to MxA-positive structures that were negative for RTN4 (and thus distinct from the standard RTN4-ER tubules), while arrows in the HA-ATL3 panels (Panel Ab) point to increased presence of RTN4- and ATL3-double positive ER “sheets” at three-way junctions. Scale bar = 20 µm. White lines in the respective merged images in Panel Aa and Ab show regions depicted in the line scans in Panel B (subpanels a and b). R values indicated in the merged images in Panel A correspond to the respective Pearson’s R coefficients (after automatic Costes’ thresholding).

**Fig. S4.**
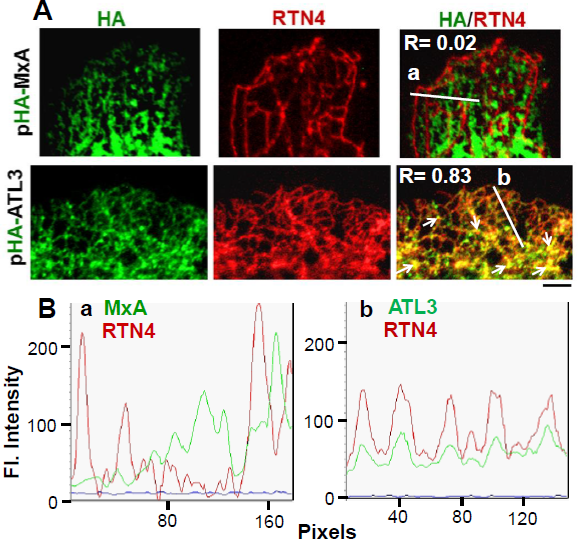
MxA-positive cytoplasmic structures in Huh7.5 cells were distinct from the standard RTN4- and ATL3-based ER (100x imaging). As indicated in the legend to Fig. S4, Huh7.5 cells were transfected with vectors for HA-tagged MxA or HA-tagged ATL3 as indicated in Panel A, the cultures (in 35 mm plates) were fixed one day later and the distribution of the HA-tag and of RTN4 sequentially evaluated using the mouse anti-HA mAb and then a goat anti-RTN4 pAb. Images were collected after placing a drop of PBS in each culture, overlaying it with a coverslip and using a 100x oil immersion objective. In Panel A, higher magnification imaging confirmed that HA-MxA did not colocalize with RTN4 tubules, while HA-ATL3 was colocalized with the RTN4-positive ER. Scale bar = 5 µm. Thin arrows point to three-way junctions that are variably positive for ATL3-HA and RTN4. White lines in the respective merged images in Panel A show regions depicted in the line scans in Panel B (subpanels a and b). R values indicated in the merged images in Panel A correspond to the respective Pearson’s R coefficients (after automatic Costes’ thresholding).

**Fig. S5.**
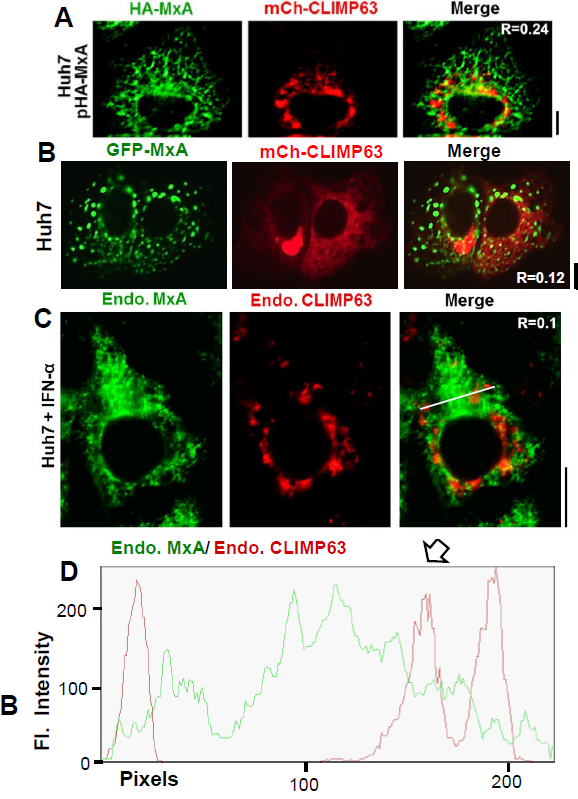
Transiently expressed HA-MxA and IFN-α-induced endogenous MxA in Huh7 cells remained distinct from CLIMP63-positive sheets of the standard ER. Panel A, Huh7 cells in 35 mm plates were transiently co-transfected with the pHA-MxA and mCherry-CLIMP63 vectors, fixed one day later and evaluated for colocalization of HA-MxA (in green using anti-MxA pAb) with mCh-CLIMP63-positive sheets of the standard ER (in red) imaged using a 40x water-immersion objective. Scale bar = 10 µm. Panel B, Huh7 cells in 35 mm plates were transiently co-transfected with the GFP-MxA and mCherry-CLIMP63 vectors, fixed one day later and evaluated for colocalization of GFP-MxA (in green) with mCh-CLIMP63-positive sheets of the standard ER (in red) imaged using a 40x water-immersion objective. Scale bar = 10 µm. Panel C, Huh7 cells were exposed to IFN-α2a (3,000 IU/ml) for 2 days, then fixed and processed for double-label immunofluorescence imaging by sequentially probing for endogenous CLIMP63 (in red using the anti-CLIMP63 mAb) and then for endogenous MxA in green (using the anti-MxA pAb) and an 100x oil-immersion objective Scale bar = 10 µm. R values indicated in the merged images in Panels A, B and C correspond to the respective Pearson’s R coefficients (after automatic Costes’ thresholding). The white line in the merged image in Panel C indicates region depicted in the line scan in Panels D.

**Fig. S6.**
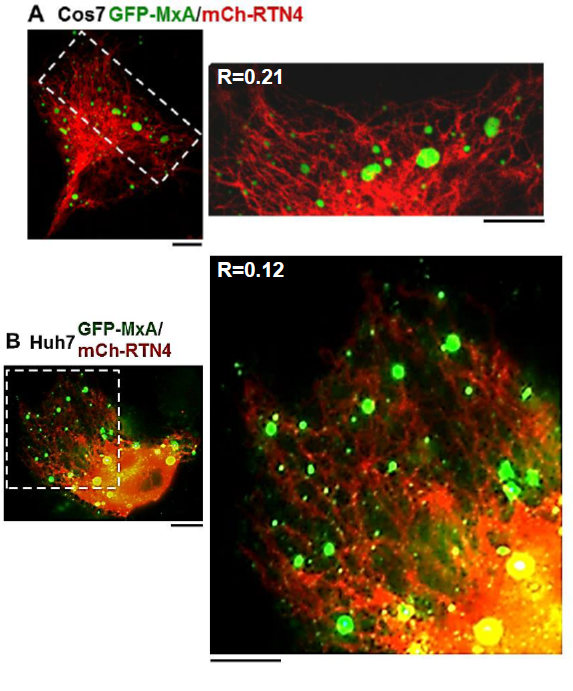
GFP-MxA positive structures in Cos7 and Huh7 cells were distinct from the standard mCh-RTN4 labelled ER. Cos7 or Huh7 cells grown in 35 mm plates as indicated were transfected with vectors for GFP-tagged MxA and mCherry-tagged RTN4 and the distribution of the respective proteins evaluated two days later following fixation and confocal 40x imaging (Panel A) or wide-field microscopy using with 100x oil imaging (Panel B). Scale bars = 10 µm in images of the entire cell (left image), and = 10 and 5 µm in the magnified insets on the right in Panels A and B respectively. Pearson’s R coefficients (with Costes’ automatic thresholding) were 0.208 and 0.119 for the data comparing GFP-MxA with mCh-RTN4 in Panels A and B respectively.

**Fig. S7.**
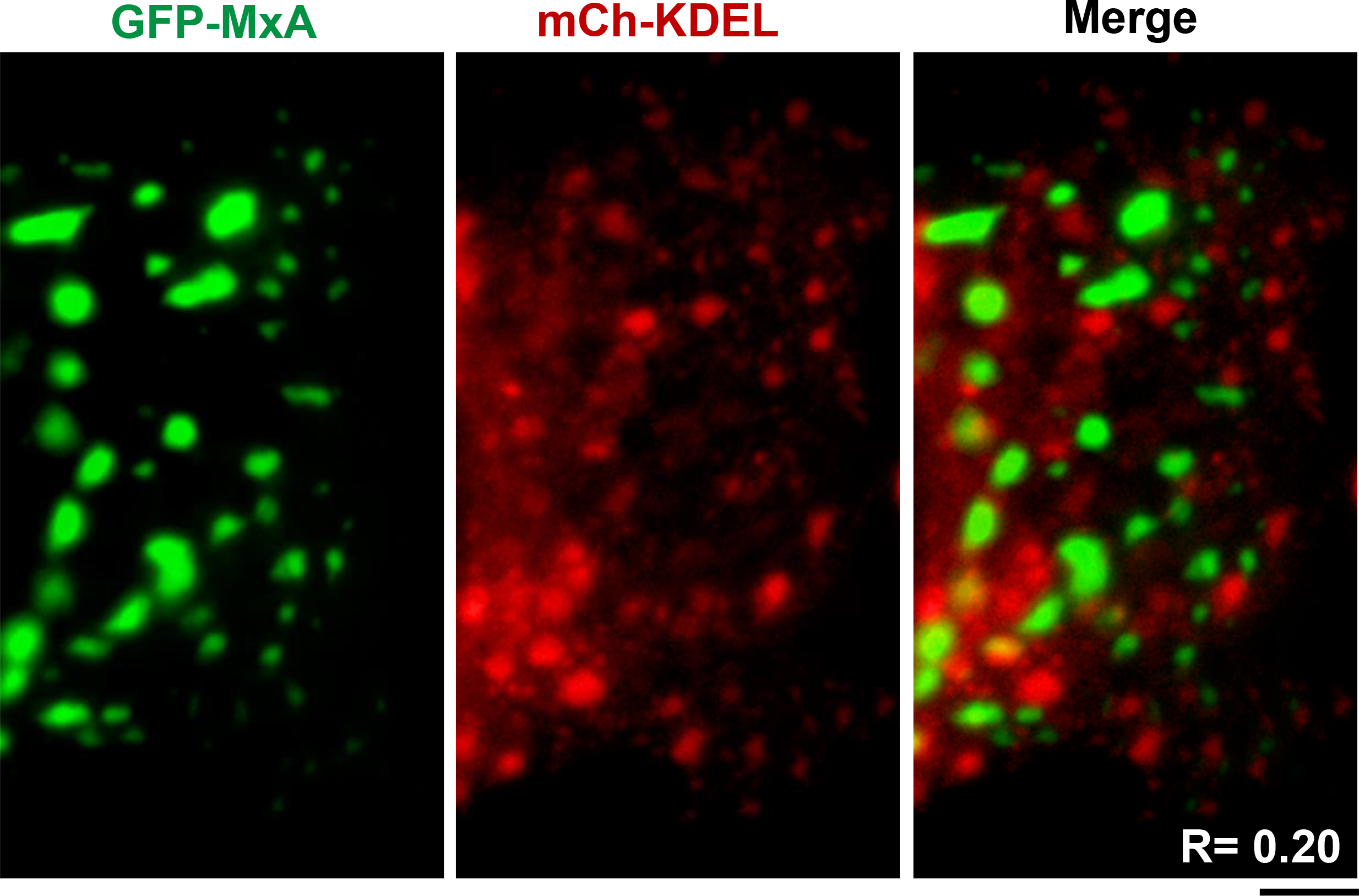
GFP-MxA positive structures in Huh7 cells were distinct from the standard mCh-KDEL labelled ER. Huh7 cells grown in 35 mm plates as indicated were transfected with vectors for GFP-MxA and mCh-KDEL and the distribution of the respective proteins evaluated two days later following fixation using wide-field microscopy using with 100x oil imaging.Scale bar = 10 µm. Pearson’s R coefficient (with Costes’ automatic thresholding) was 0.20 for the data comparing GFP-MxA with mCh-KDEL.

**Fig. S8.**
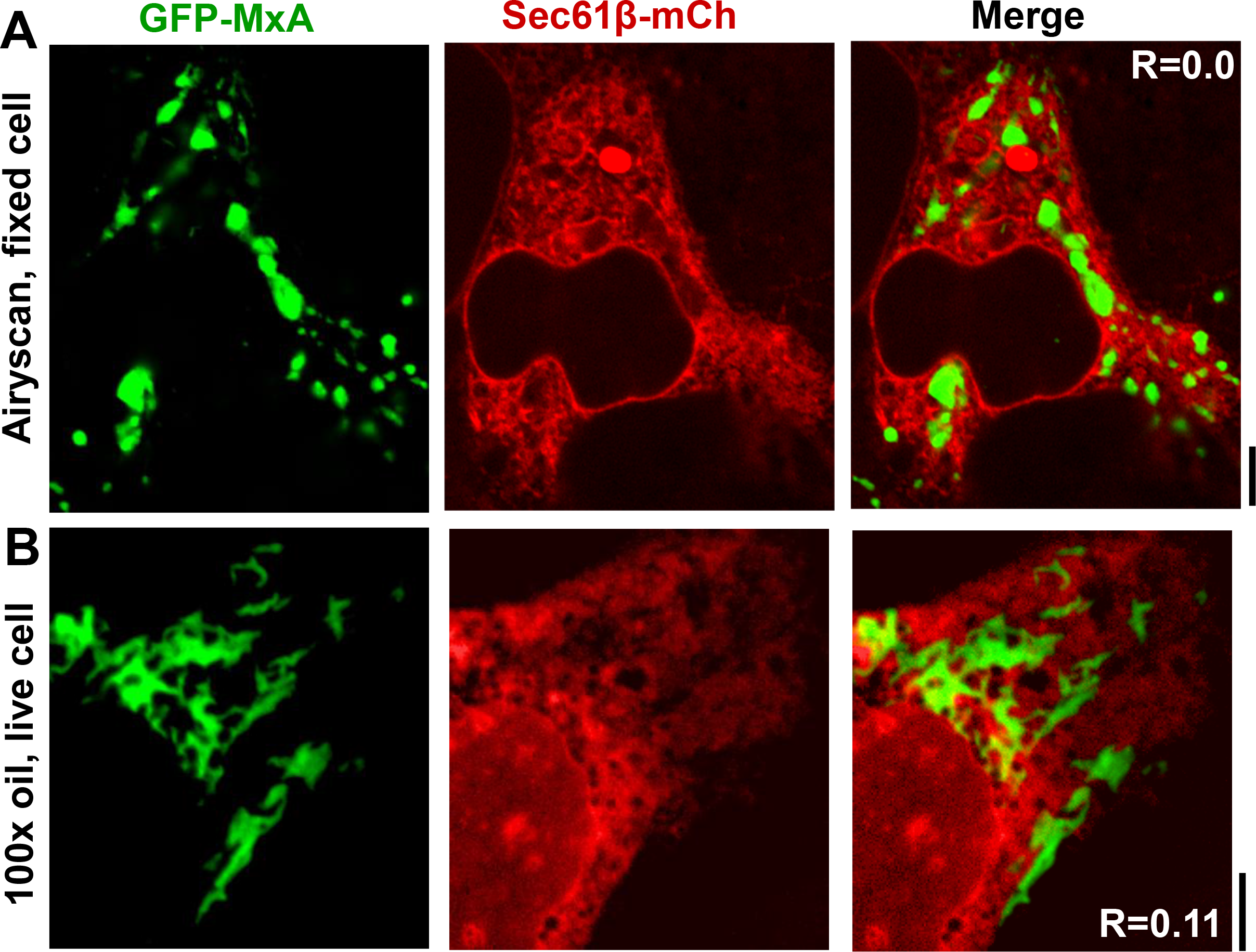
GFP-MxA positive structures in Huh7 cells were distinct from mCh-Sec61β-tagged standard ER. Huh7 cells in 35 mm plates were transiently co-transfected with the vectors for GFP-MxA and mCh-Sec61β, fixed 2 days later and evaluated for colocalization of the respective proteins using a high-resolution Zeiss confocal Airyscan system (Panel A) in an experiment carried out contemporaneously with initiating the CLEM analyses in Fig. S9. Panel B shows data from an independent live-cell imaging experiment. Scale bars = 5 µm. R values indicated in the merged images in Panels A and B (= 0.0 and 0.11 respectively) correspond to the respective Pearson’s R coefficients (after automatic Costes’ thresholding).

**Fig. S9.**
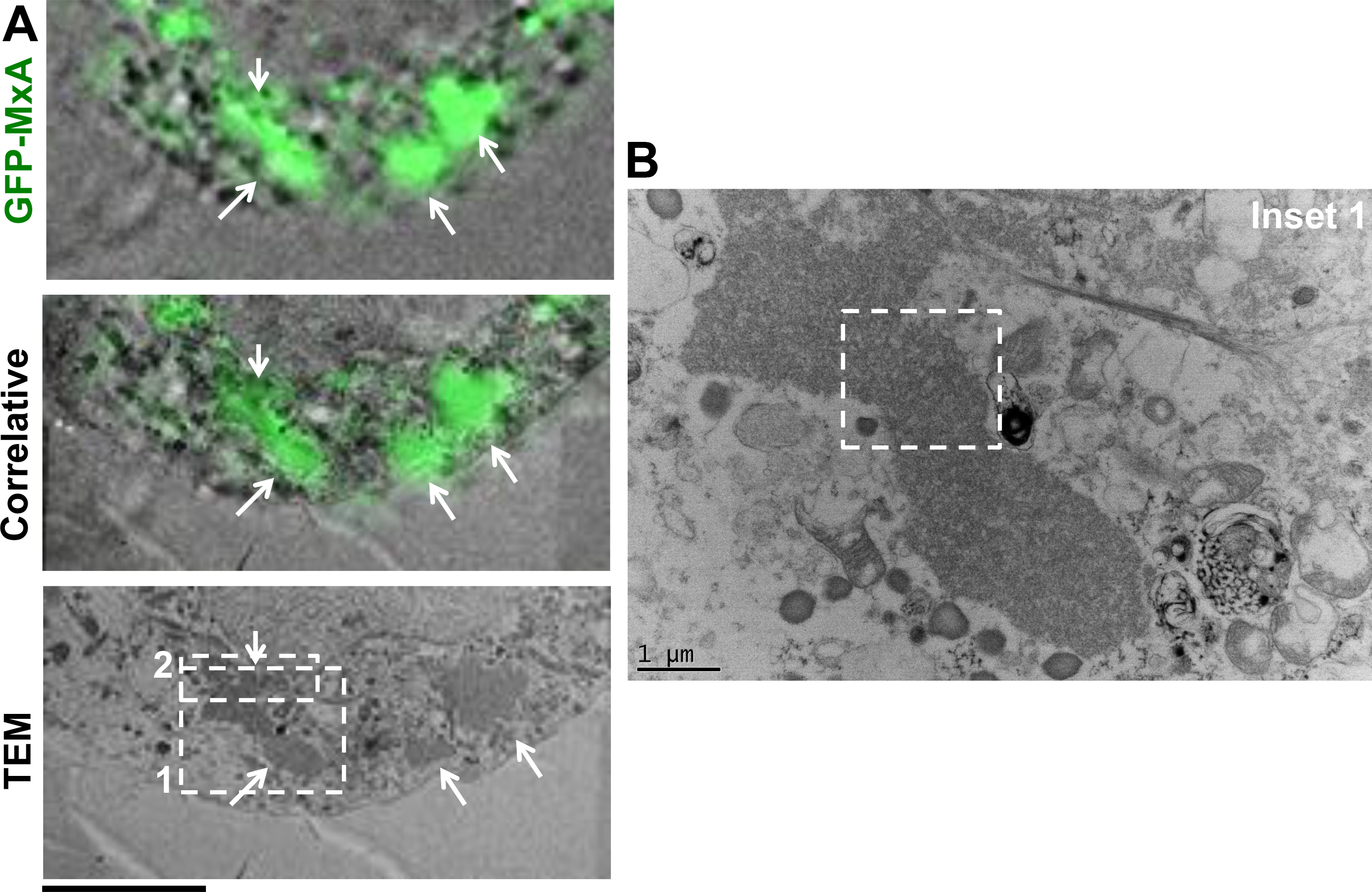
Correlated light and electron microscopic (CLEM) analyses of GFP-MxA structures in Huh7 cells. Huh7 cells plated sparsely in 35 mm gridded coverslip plates were transiently co-transfected with the pGFP-MxA. Two days later the cultures were fixed with 4% paraformaldehyde for 1 hr at 4°. Confocal imaging was carried out using a tiling protocol to identify the location of specific cells with GFP-MxA structures on the marked grid. The cultures were then further fixed, embedded and the previously identified grid locations used for serial thin-section EM (TEM). The tiled light microscopy data were correlated with the tiled EM data to identify the ultrastructure of the GFP-fluorescent structures (arrows in Panel A). Scale bar = 5 µm. Panel B shows a higher magnification image of the inset 1 indicated in bottom. Scale bar = 1 µm. Higher magnification TEM images of inset 2 in Panel A are shown in Fig. 3B, and of the boxed inset in Panel B in Fig. 3A.

**Fig. S10.**
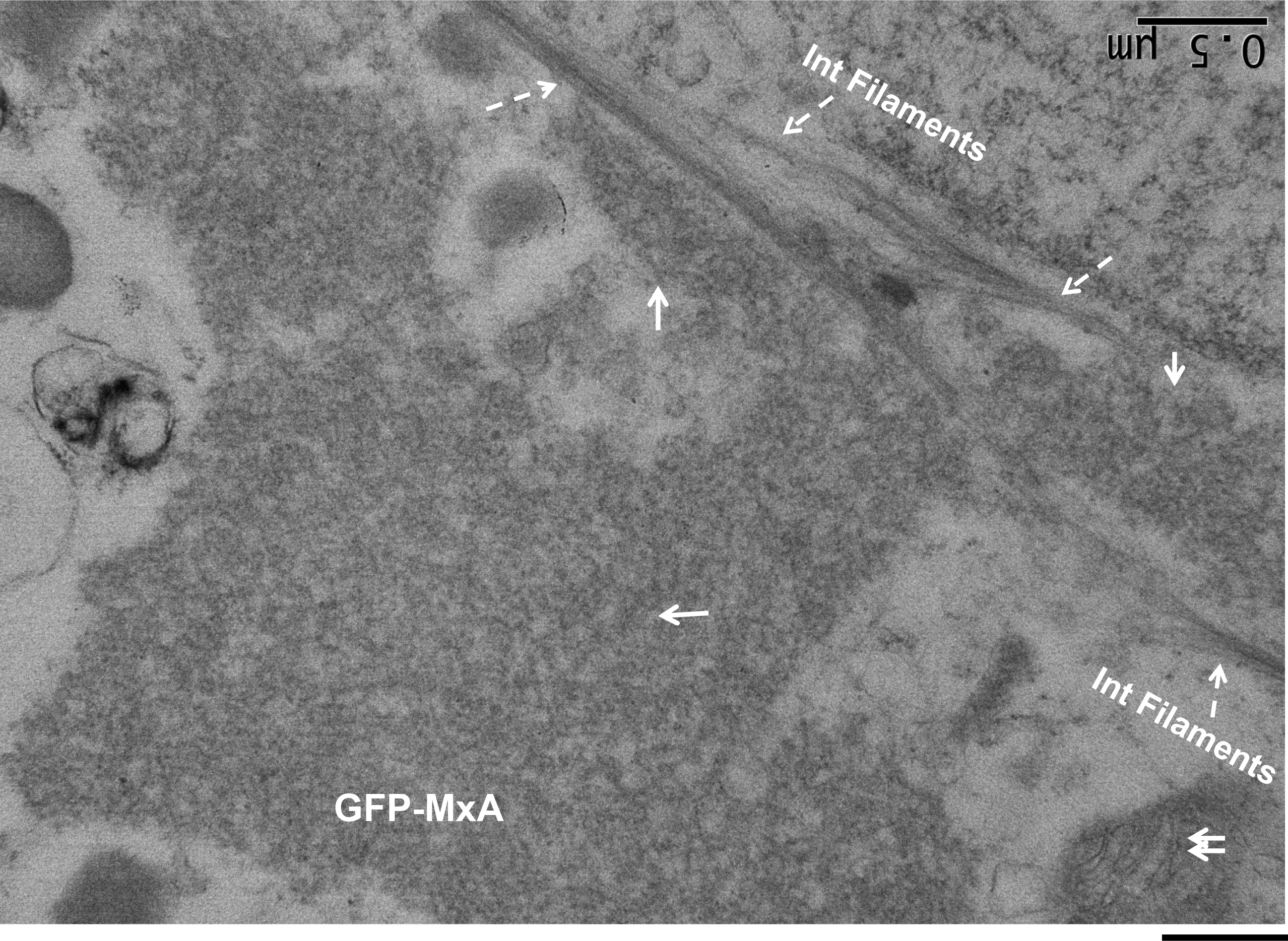
Correlated light and electron microscopic (CLEM) analyses of GFP-MxA structures in Huh7 cells – association with intermediate filaments. This figure illustrates an higher magnification TEM image of the upper left portion of Fig. S9A, Panel B. Solid arrows point to GFP-MxA membrane-less structures, while broken arrows point to intermediate filaments. The double arrow in the lower right corner point to cytoplasmic membranes observed within the same image (as a positive control verifying that the GFP-MxA structures lacked an external limiting membrane as observed in the same image). Scale bar = 500 nm.

**Fig. S11.**
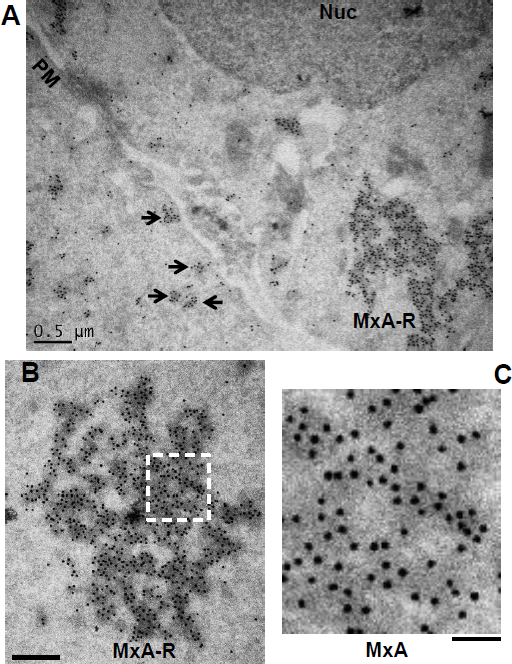
Identification of cytoplasmic MxA structures of varied morphology by immuno-EM. HA-MxA expressing HEK293T cells (as in Fig. 1A) were prepared for immuno-EM analyses of MxA-positive structures using rabbit pAb to MxA and 15 nm Protein A-gold as the secondary label. Panel A, shows strong labeling of small and variably-sized MxA bodies (arrows) as well as larger MxA reticular structures (MxA-R). PM, plasma membrane, Nuc, nucleus. Panels B shows a different example of MxA-R, while Panel C shows the boxed inset in Panel B at a higher magnification. Scale bars = 500 nm in Panels A and B, and 200 nm in Panel C.

**Fig. S12.**
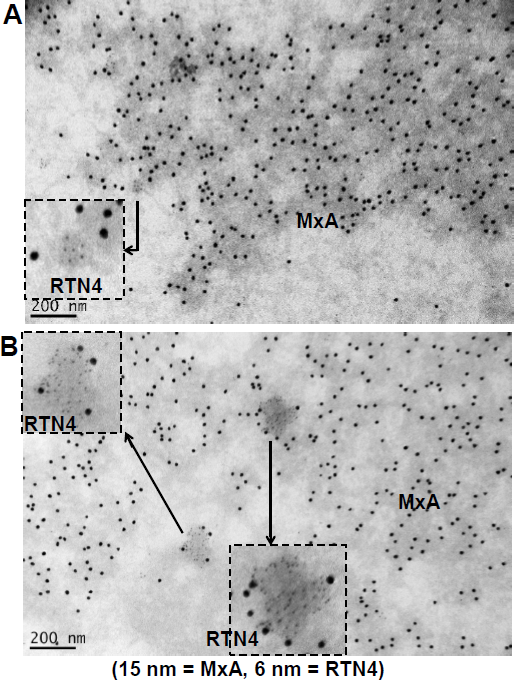
Double-labeled immuno-EM showing that the MxA-reticular structures were largely negative for RTN4. Double-labeled immuno-EM studies were carried out on HEK293T cells expressing HA-MxA (as in Fig. 1A) by labeling MxA with 15 nm colloidal gold and RTN4 with 6 nm colloidal gold. Panel A, the MxA-reticulum observed was exclusively MxA-positive; a few RTN4-ER structures (cut in cross-section) were present adjacent to the MxA-reticulum (see boxed inset). Panel B, the standard RTN4-ER was largely devoid of MxA. However, boxed insets show higher magnification of two examples of RTN4-positive ER structures which with some evidence for peripheral MxA proximity. Critically, Panel B demonstrates the validity of the double-label immuno-EM method by evidencing MxA and RTN4-postive structures labeled with different-sized gold particles in the same image. Scale bars = 200 nm.

**Fig. S13.**
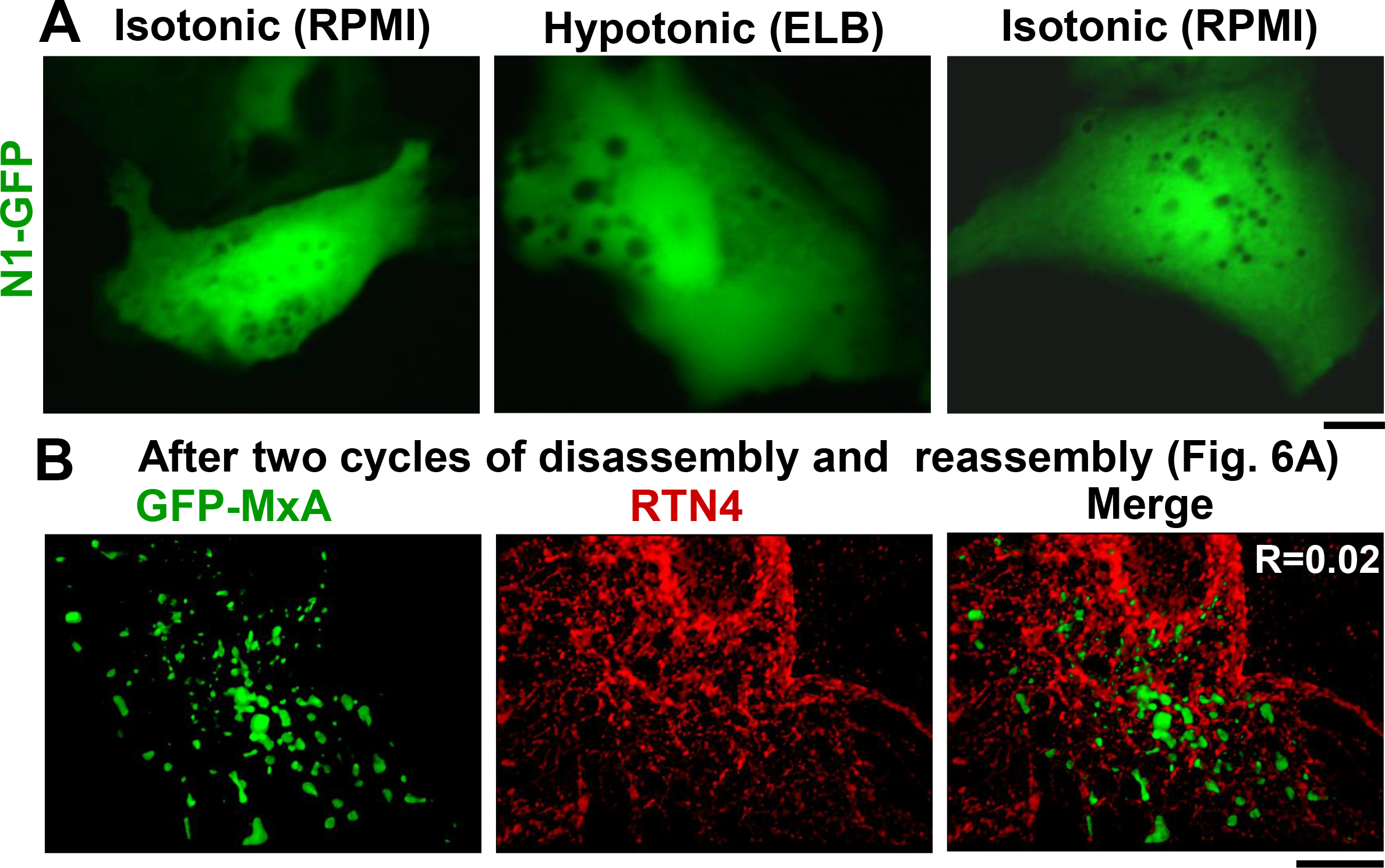
Lack of condensate formation by N1-GFP and verification that reassembled GFP-MxA condensates were distinct from the RTN4-ER. Panel A, Huh7 cells transiently expressing the N1-GFP tag alone were imaged in isotonic medium (RPMI), after switching to hypotonic buffer (ELB) for approximately 10 min, and then after switching back to isotonic medium (RPMI) for approximately 10 min. Panel B. Cultures in the cycled through two rounds of disassembly and reassembly shown in Fig. 8A were fixed, and evaluated for colocalization of the reassembled GFP-MxA structures with RTN4 (in red by immunofluorescence). R value indicated in the merged image (R = 0.02) corresponds to the Pearson’s R coefficient (after automatic Costes’ thresholding).

**Fig. S14.**
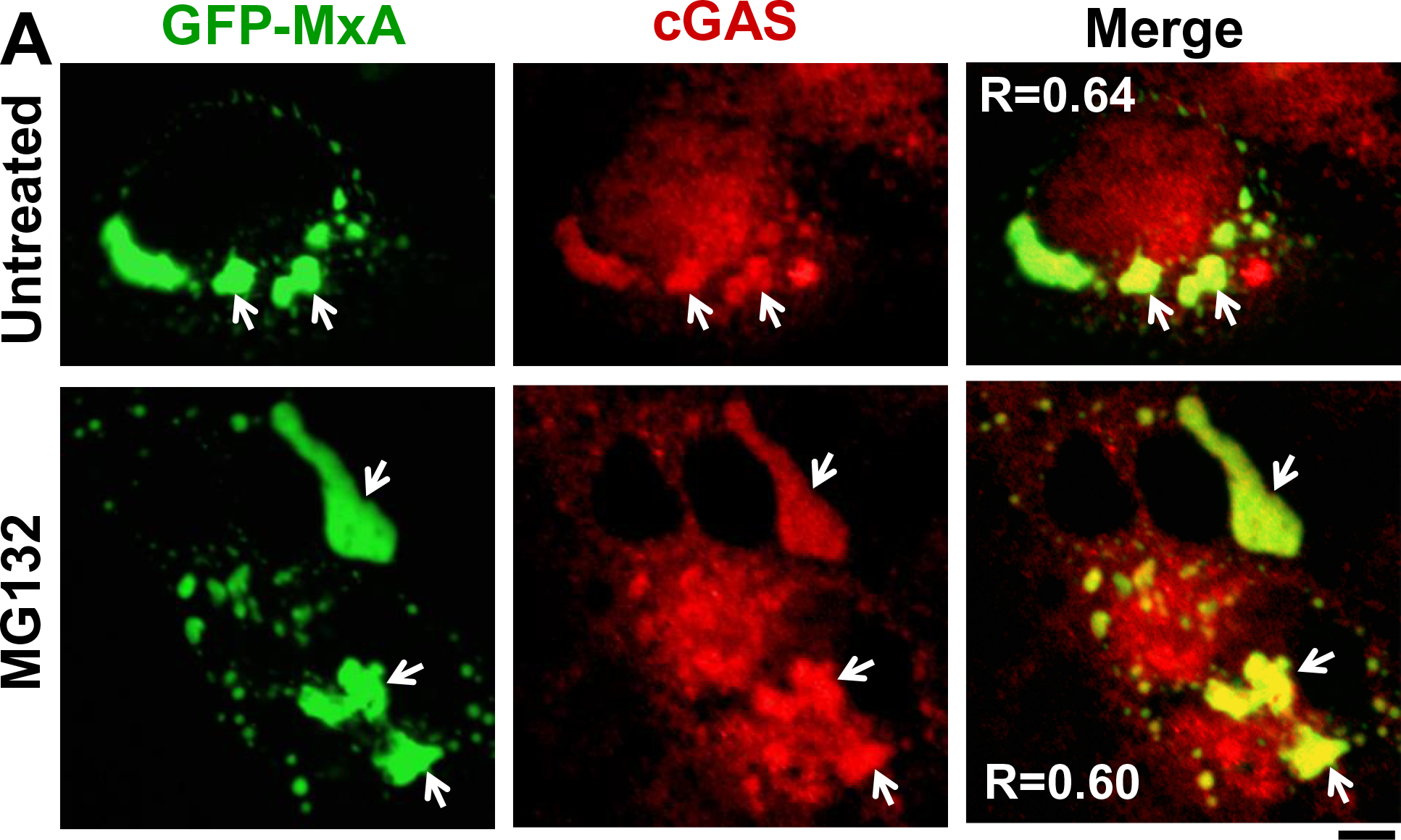
Inclusion of cGAS in GFP-MxA condensates. GFP-MxA expressing cultures 2 days after transfection were either left untreated or treated with the proteasome inhibitor MG132 (40 µM) for 4 hr. The cultures were fixed, permeabilized, and cGAS detected using immunofluorescence (in red) and compared to GFP-MxA (in green). R is the Pearson’s correlation coefficient after automatic Costes’ thresholding. Scale bar = 20 µm.

## Legends to Supplementary Movies

**Movie S1. Homotypic association of variably-sized GFP-MxA bodies.** This time-lapse movie corresponds to the experiment and the specific cell illustrated in Fig. 5, Panel A.

**Movie S2. Oscillatory movement of GFP-MxA filaments and reticulum.** This time-lapse movie corresponds to the experiment illustrated in Fig. 7, Panel B (Huh7 cell treated with c-PTIO, 300 µM, 7 hr).

**Movie S3. Oscillatory movement of GFP-MxA filaments and reticulum.** This time-lapse movie corresponds to the experiment and the specific cell illustrated in Fig. 7, Panel C (Huh7 cell treated with dynasore, 20 nM, for 4 days).

**Movie S4. Rapid disassembly of GFP-MxA condensates upon exposing Huh7 cells to hypotonic buffer and reassembly upon shifting to isotonic buffer.** The same cell is illustrated through both the disassembly and reassembly cycles. This time-lapse movie corresponds to an experiment similar to that illustrated in Fig. 8, Panel A, cycle 1 (Change from isotonic RPMI to hypotonic ELB and then back to isotonic buffer).

**Movie S5. Rapid reassembly of GFP-MxA condensates upon exposing Huh7 cells to isotonic buffer.** This time-lapse movie corresponds to the experiment and the specific cell illustrated in Fig. 8, Panels C and D, cell (c) (change from hypotonic ELB to isotonic RPMI).

